# The polyadenosine RNA binding protein Nab2 regulates alternative splicing and intron retention during *Drosophila melanogaster* brain development

**DOI:** 10.1101/2024.05.17.594635

**Authors:** Seth M. Kelly, Allison Paschack, Gargi Mishra, Jinsoo Ahn, Emma Smith, Katie Haynes, Katherine Shelmidine, Kichoon Lee, Andre Yazhbin, Young Ji

## Abstract

The regulation of cell-specific gene expression patterns requires the coordinated actions of hundreds of proteins, including transcription factors, processing enzymes, and many RNA binding proteins (RBPs). RBPs often become associated with an mRNA immediately after its production and can coordinate processing and quality control steps. Since RBPs can regulate multiple post-transcriptional processing steps, mutations within RBP-encoding genes often lead to pleiotropic effects that alter the physiology of multiple cell types. Thus, identifying how an RBP functions during RNA processing can provide a better understanding of both tissue physiology and mechanisms of disease.

In the current study, we investigated how the *Drosophila* RNA binding protein Nab2, an evolutionary conserved ortholog of human ZC3H14, coordinates mRNA splicing and polyadenylation. ZC3H14 loss in human patients has previously been linked to alterations in nervous system function and disease. Both fly Nab2 and vertebrate ZC3H14 bind to polyadenosine RNA and have been implicated in the control of poly(A) tail length. Here we show that *Nab2* loss from developing fly brain tissue also causes widespread changes in alternative splicing and intron retention. Retained introns in Nab2 null larval brains are most often first introns, are surrounded by adenosine-rich sequences, and are often associated with extended poly(A) tails, suggesting that Nab2 coordinates the splicing and polyadenylation of target transcripts. Alterations in splicing patterns ultimately lead to changes in protein abundance in Nab2 null developing brains. Overall, these studies highlight the importance of Nab2 and ZC3H14 in the post-transcriptional regulation of gene expression and provide critical insight into how this family of RBPs coordinates RNA processing during brain development.

## Introduction

During organismal development, post-transcriptional mechanisms regulate gene expression to gradually refine patterns of protein production and contribute to the formation of specific cell types (Raj and Blencowe 2015; Lennox et al. 2018; Weyn-Vanhentenryck et al. 2018; Lee et al. 2024). Splicing is one such processing step that is tightly regulated and crucial for establishing specific patterns of isoform expression. For example, within developing *Drosophila* larval sensory neurons, the alternative splicing of *Dscam1* transcripts promotes self-avoidance of sensory neuron dendrites during dendrite outgrowth (Wojtowicz et al. 2004; Matthews et al. 2007; Soba et al. 2007). Alterations in typical *Dscam1* splicing patterns also cause axonal extension defects in the developing *Drosophila* mushroom body, a brain region required for olfactory learning and memory in flies (Wang et al. 2002; Zhan et al. 2004; Hattori et al. 2007; Lin 2023). Multiple processing events occur during and after transcription to regulate the final abundance and sequence of mature mRNAs.

Splicing is facilitated by a multi-subunit complex called the spliceosome which is made from dozens of proteins and RNAs (Will and Luhrmann 2011; Matera and Wang 2014). The spliceosome consists of five small nuclear ribonucleoprotein complexes (snRNPs), each of which contain a U-rich RNA transcript (Matera and Wang 2014). The U-rich snRNA transcripts of the major spliceosome (U1, U2, U4, U5, and U6) are transcribed in the nucleus. Most snRNAs (except U6) are exported to the cytoplasm and assembled with a ring of Sm proteins (SmB, SmD1, SmD3, SmE, SmF, and SmG) in a stepwise manner by the pICln and Survival of Motor Neuron (Smn) complexes (Patel and Bellini 2008; Matera and Wang 2014; Neuenkirchen et al. 2015). The fully assembled snRNPs are then imported into the nucleus where they appear to be stored in locations called nuclear speckles (Spector and Lamond 2011; Matera and Wang 2014; Faber et al. 2022). Eventually, snRNPs assemble at pre-mRNA splice sites and facilitate the removal of introns. In most cases, soon after it is transcribed, the pre-mRNA 5’-splice site (5’ss) is recognized by the U1 snRNP. The intron branchpoint sequence within the intron is then recognized by the branchpoint binding protein and the U2 snRNP (Pomeranz Krummel et al. 2009; Matera and Wang 2014; van der Feltz and Hoskins 2019; Zhang et al. 2020b). Following the recruitment of the U4/U6.U5 tri-snRNP and a series of rearrangements, the branchpoint, 5’ss, and 3’ss are brought into closer proximity and a splicing-competent (i.e. activated) spliceosome is formed (Will and Luhrmann 2011; Matera and Wang 2014; Didychuk et al. 2018; Kastner et al. 2019). Eventually, the 5’ss is attached to the 3’ss and the intervening intron lariat is released and degraded (Matera and Wang 2014). Throughout the splicing process, multiple RNA helicases rearrange the spliceosome into its active conformation and disassemble the spliceosome following lariat release (Cordin and Beggs 2013).

The efficiency of splice site usage and alternative splicing patterns are often determined by RNA binding proteins (RBPs) bound to RNA sequences near splice sites (Fu and Ares 2014; Raj and Blencowe 2015). Binding of RBPs to RNA sequences near the splice site may lead to the inclusion or exclusion of nearby exons and result in tissue-specific patterns of alternative splicing. For example, the mammalian family of Rbfox proteins (including Rbfox1, Rbfox2, and Rbfox3) are preferentially expressed in the nervous system, bind to intronic sequences upstream or downstream of regulated exons, and can then decrease or increase, respectively, the likelihood that the exon will be included in the final processed mRNA transcript (Yeo et al. 2009; Lovci et al. 2013; Fu and Ares 2014; Misra et al. 2015). High throughput sequencing approaches have also demonstrated changes in exon inclusion patterns during pre- and post-natal brain development that are correlated with changes in the abundances of several RNA binding proteins, including Rbfox, Ptbp, Nova, and Mbnl family members (Raj and Blencowe 2015; Weyn-Vanhentenryck et al. 2018). These studies and others have demonstrated that increased expression of some proteins (such as Rbfox and Nova) and the decreased expression of others (such as Ptbp family members) during cortical development is correlated with increased levels of neuronal differentiation (Charizanis et al. 2012; Licatalosi et al. 2012; Weyn-Vanhentenryck et al. 2014; Raj and Blencowe 2015; Weyn-Vanhentenryck et al. 2018), suggesting a critical role for RBPs in organismal development.

Splicing, capping, and polyadenylation are often completed simultaneously and the effectiveness of one step can influence the others (Misra et al. 2015; Herzel et al. 2018; Zhang et al. 2023). RNA binding proteins involved in multiple aspects of RNA metabolism are uniquely positioned to help coordinate these processing events and ensure the correct expression of targeted transcripts. For example, vertebrate ZC3H14 (also referred to as mSUT2) is an evolutionarily conserved RBP that has been implicated in poly(A) tail length control, transcriptional elongation, and mRNA splicing (Kelly et al. 2007; Pomeranz Krummel et al. 2009; Pak et al. 2011; Kelly et al. 2014; Soucek et al. 2016; Wigington et al. 2016; Rha et al. 2017; Morris and Corbett 2018). Mutations in *ZC3H14* have been linked to human intellectual disability (Pak et al. 2011), liver cancer (Li et al. 2019), and tauopathy (Guthrie et al. 2009; Guthrie et al. 2011). ZC3H14 binds to poly(A) RNA (Kelly et al. 2007) via C-terminal tandem Cys-Cys-Cys-His (CCCH) zinc fingers and has been immunoprecipitated with components of the transcriptional elongation THO complex (Morris and Corbett 2018). ZC3H14 is localized within nuclear speckles, the proposed storage sites for multiple spliceosome components (Pomeranz Krummel et al. 2009; Spector and Lamond 2011), and loss of ZC3H14 also causes the accumulation of unspliced pre-mRNA in the nucleus (Wigington et al. 2016) and extended poly(A) tails (Pak et al. 2011; Bienkowski et al. 2017). The yeast and fruit fly orthologs of ZC3H14, called Nab2 (and in some studies, dNab2), have also been implicated in both poly(A) tail length control and pre-mRNA splicing (Kelly et al. 2014; Soucek et al. 2016; Jalloh et al. 2023; Lancaster et al. 2024). Yeast Nab2 genetically and physically interacts with components of the spliceosome (Soucek et al. 2016) and is necessary for correct splicing of specific transcripts (Soucek et al. 2016; Wigington et al. 2016). Like their vertebrate counterpart, both yeast and *Drosophila* Nab2 localize to the nucleus, regulate bulk poly(A) tail length, and bind directly to polyadenosine RNA via C-terminal CCCH zinc fingers (Anderson et al. 1993; Hector et al. 2002; Pak et al. 2011; Kelly et al. 2014). Importantly, Nab2 null flies are mostly inviable but can be rescued by selective expression of human ZC3H14 within neurons, suggesting that fly Nab2 and human ZC3H14 perform similar molecular functions within the nervous system (Pak et al. 2011; Kelly et al. 2014). Finally, more recent work has also demonstrated that adult flies lacking Nab2 have atypical splicing patterns and m^6^A hypermethylation of transcripts encoding the sex-lethal (sxl) RBP (Jalloh et al. 2023), implicating Nab2 in control of RNA processing and modification. Given the co-transcriptional nature of many processing events such as splicing and polyadenylation, the exact step of RNA processing where Nab2 functions has been difficult to unravel. The implications of ZC3H14 loss in human patients has likewise been difficult to ascertain. Importantly, reported phenotypes observed in adult ZC3H14 null mice or Nab2 null adult flies also might be confounded by defects at earlier stages, necessitating a more thorough analysis of ZC3H14/Nab2 loss across developmental time periods. In the current study, we have investigated the genome-wide consequences of *Drosophila melanogaster* Nab2 loss on both mRNA splicing and polyadenylation during larval brain development. We show here that Nab2 genetically interacts with core splicing components and prevents intron retention within selected mRNA transcripts. Our findings suggest that Nab2 loss from developing larval brain tissue caused reproducible changes in the expression levels of hundreds of genes, many of which are unique when compared to changes that occur in Nab2 null adult flies (Jalloh et al. 2023). Intron retention within several transcripts in Nab2 null larval brain tissue also resulted in protein production alterations during brain development. Although previous studies have demonstrated a modest global increase in poly(A) tail lengths when ZC3H14 or Nab2 is missing (Pak et al. 2011; Kelly et al. 2014; Bienkowski et al. 2017; Rha et al. 2017), long-read RNA sequencing from larval brain tissue demonstrated that Nab2 is not required for poly(A) tail length control on most mRNA transcripts during larval brain development. Instead, those transcripts that require Nab2 for correct splicing appear to possess significantly longer poly(A) tails in the absence of Nab2, providing novel insight into the role of Nab2 in coordinating the splicing and polyadenylation of these target mRNAs. Together, this work advances our understanding of how RNA processing events are coordinated by RNA binding proteins during brain development and may provide for a better understanding of molecular defects that occur in human patients with ZC3H14 mutations.

## Results

### *Drosophila Nab2* genetically interacts with genes encoding components of the spliceosome

Previous data suggests that *S. cerevisiae* Nab2 associates with multiple RNA processing proteins, including several splicing regulators (Soucek et al. 2016), RNA exosome factors (Soucek et al. 2016), and 3’-end cleavage and polyadenylation complex members (Kelly et al. 2010). However, yeast Nab2 has mainly been implicated in the control of polyadenylation and RNA export from the nucleus (Hector et al. 2002; Marfatia et al. 2003). We hypothesized that in other species where alternative splicing is more prevalent, Nab2 and its human ortholog ZC3H14, might play a more prevalent role in regulating alternative splicing. To test functional interactions between *Nab2* and genes encoding components of the spliceosome, we investigated whether RNA interference (RNAi)-mediated knockdown of splicing protein expression in the fly eye could dominantly modify a rough eye phenotype caused by Nab2 overexpression (Pak et al. 2011). As shown in Figure 1, overexpression of Nab2 in the fly eye caused a slightly smaller and more disorganized adult eye. By comparison, flies containing one copy of the *ninaE.GMR-GAL4* driver had typical eyes with well-ordered ommatidia. When expression of the Sm proteins SmB, SmD3, SmE, or SmF was decreased in eyes also overexpressing Nab2, the rough eye phenotype was substantially enhanced, and eyes became more disorganized and marked by black necrotic tissue in the posterior half of the adult eye (Fig 1A, posterior is to the right of image and marked with a P). Consistent enhancement of eye roughness was often observed using multiple independent RNAi lines (Supplemental Table 1) targeting the same gene, suggesting that these results were not due to off-target effects of individual RNAi lines. Importantly, decreasing expression of individual Sm proteins alone without altering Nab2 levels in the fly eye caused no change in eye morphology (bottom row, Fig 1A).

**Fig 1:**
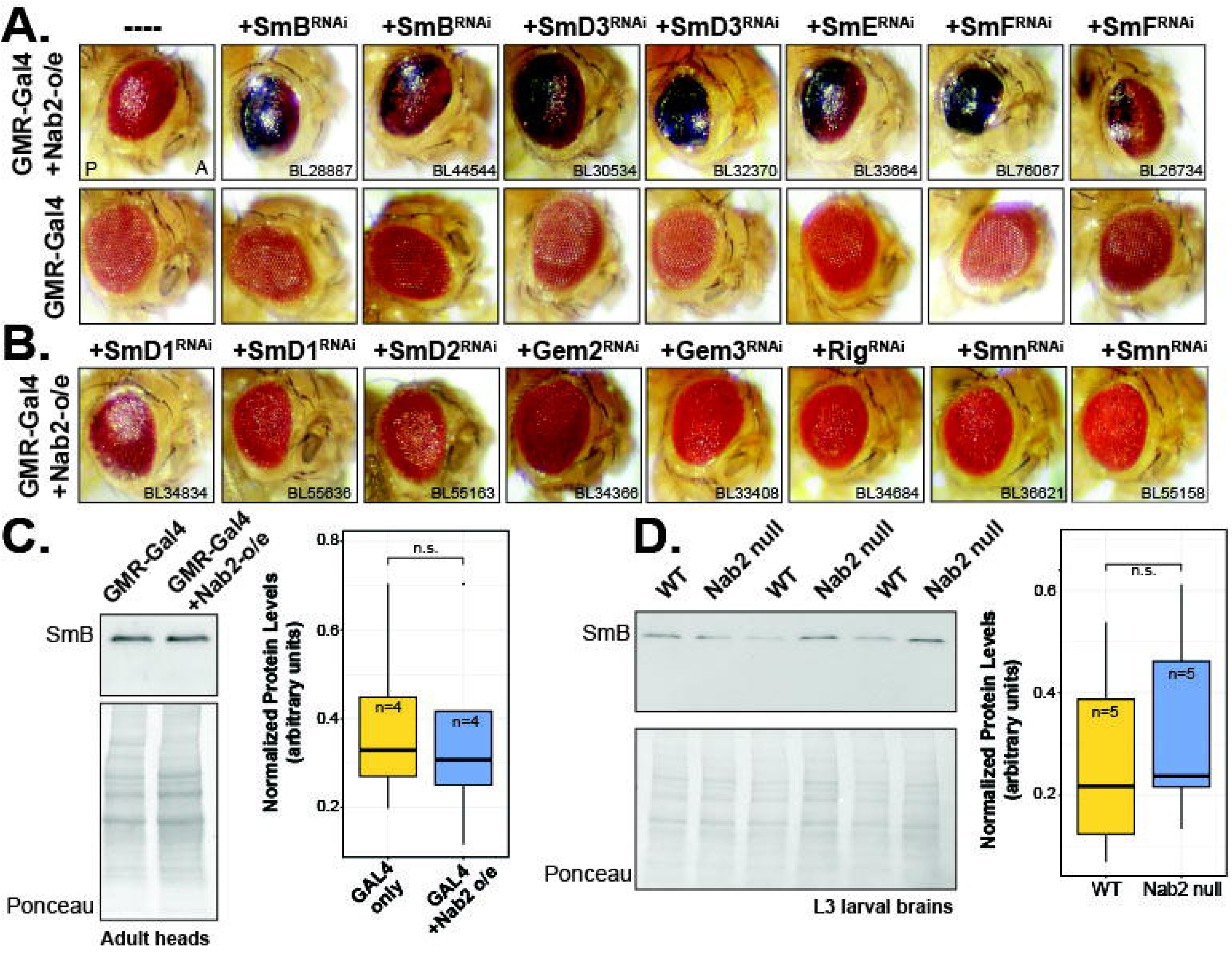
Loss of Sm protein expression genetically modifies *Drosophila melanogaster* eye morphology phenotypes caused by Nab2 overexpression. (A) Overexpression of Nab2 in the fly eye (genotype: *ninaE.GMR-GAL4/+; Nab2^EP3716^/+*) caused eye disorganization. Control flies containing *GAL4* only (genotype: *ninaE.GMR-GAL4/+*) did not have eye morphology defects. Decreased expression of SmB, SmD3, SmE, or SmF in fly eyes by RNA interference (RNAi) enhanced Nab2-induced eye roughness (top row, genotypes: *ninaE.GMR-GAL4/+; Nab2^EP3716^/UAS-Sm Protein-RNAi*). All *UAS-Sm-RNAi* lines were combined with *ninaE.GMR-GAL4* (bottom row) to control for morphology defects caused by splicing protein knockdown. The stock number of the RNAi line used is shown. (B) Decreased expression of SmD1, SmD2, or components of the Gemin/Smn complex did not enhance Nab2 overexpression induced fly eye morphology defects. (C) Nab2 overexpression is not sufficient to alter levels of SmB protein in fly heads. Soluble protein from control (*ninaE.GMR-GAL4/+*) fly heads and fly heads overexpressing Nab2 in fly eyes (*ninaE.GMR-GAL4/+; Nab2^EP3716^/+*) was used to analyze levels of SmB protein by immunoblot. Box and whisker plots show the abundance of SmB in each condition normalized to Ponceau staining from four biological replicates. No significant difference was observed by two-sample t-test (t(6) = 0.18479, p > 0.859). (D) Nab2 is not necessary for control of SmB levels in *Drosophila* larval brain tissue. SmB protein was detected in protein samples from control (*Nab2^pex41^/Nab2^pex41^*) or Nab2 homozygous null (*Nab2^ex3^/Nab2^ex3^*) L3 larval brain lysates (n=3). No significant difference in SmB levels was observed by two-sample t-test (t(8) = -0.52507, p = 0.6138).

Sm proteins assemble into specific subcomplexes first and are then loaded onto U-rich snRNAs by the Survival of Motor Neuron (Smn) complex before being reimported into the nucleus (Matera and Wang 2014; Neuenkirchen et al. 2015). Interestingly, our data suggest that Nab2 functionally interacts with SmB/SmD3 and SmE/SmF/snRNPG complex but not with the SmD1/SmD2 subcomplex, as decreasing expression of SmD1 or SmD2 in the fly eye did not enhance the rough eye phenotype caused by Nab2 overexpresson (Fig 1B). Likewise, reduced expression of Smn complex proteins, such as Gemin 2 (Gem2), Gemin 3 (Gem3), Gemin 5 (called Rig in *D. melanogaster*), or Smn did not enhance the rough eye phenotype caused when Nab2 is overexpressed in the fly eye (Fig 1B).

One straight-forward molecular explanation for the dominant enhancement of the rough eye phenotype seen when Sm protein levels were reduced is that Nab2 regulates the expression of specific Sm proteins, such as SmB. Overexpression of Nab2 in the fly eye could then cause alterations that are further enhanced by Sm protein knockdown. To test this idea, we analyzed the level of SmB protein in adult fly heads where Nab2 was overexpressed only in fly eyes. As shown in Fig 1C, the level of SmB protein present in flies overexpressing Nab2 (genotype: *ninaE.GMR-GAL4>Nab2^EP3716^*) was not significantly different than in control flies (genotype: *ninaE.GMR-GAL4*, p > 0.05). To look more holistically at whether Nab2 regulates SmB protein levels, we also analyzed the level of SmB protein in *Nab2^ex3^/Nab2^ex3^* (hereafter called Nab2 null) larval brain tissue (Fig 1D). *Nab2^ex3^* is an RNA and protein null allele of Nab2 caused by imprecise excision of the P-element found in *Nab2^EP3716^* (Pak et al. 2011). The loss of Nab2 from larval brain tissue did not significantly alter the level of SmB protein compared to control larval brain tissue (genotype: *Nab2^pex41^/Nab2^pex41^*, where EP3716 was removed from the *Nab2* locus precisely, p > 0.05, (Pak et al. 2011)).

We next investigated whether *Nab2* genetically interacts with other splicing components, including members of the U1 snRNP (Fig 2B), the U2 snRNP, the U4/U5.U6 tri-snRNP (Fig 2C), and several RNA helicases (Fig 2D) required for remodeling of the spliceosome between splicing steps (Matera and Wang 2014). We again utilized the eye-specific *ninaE.GMR-Gal4* line to increase expression of Nab2 and simultaneously decrease expression of selected splicing components with UAS-driven RNAi constructs (Perkins et al. 2015). Similar to the image shown previously in Figure 1, overexpression of Nab2 in the fly eye (genotype: *ninaE.GMR-Gal4 + Nab2^EP3716^*) caused disorganization of eye facets (Fig 2A). Nab2 induced eye roughness was not genetically modified when the levels of U1 snRNP components, including Prp40, snf, snRNP-U1-70K, and snRNP-U1-C (Fig 2B and Supplemental Table 1), were reduced. However, eye disorganization was enhanced when levels of U2 snRNP components were reduced in combination with Nab2 overexpression (Fig 2C). Decreased levels of SF3A1 and Spx (the fly ortholog of SF3B4 (Herold et al. 2009)) caused significant enhancement of eye roughness when Nab2 was simultaneously overexpressed in the fly eye. Fly eyes were smaller in size and mostly consisted of black necrotic tissue, especially in the posterior half of the eye. Importantly, SF3A1 knockdown on its own caused very minimal changes to eye morphology that were barely detectable and Spx knockdown on its own caused no discernable changes to eye morphology (Fig 2C and Supplemental Table 1). Knockdown of U2AF50 (known as Mud2 in *S. cerevisiae*), also caused dominant enhancement of the Nab2 rough eye phenotype but did not have an effect on its own (Fig 2C). Reduction in the levels of other U2 snRNP components, including U2A, SF3B2, SF3B3, and noi did not significantly alter the eye roughness phenotype caused by Nab2 overexpression (Supplemental Table 1). Other U2 snRNP components, such as SF3B1, were not testable in this experiment because they caused lethality or eye roughness when their expression was decreased with *ninaE.GMR-GAL4* alone (Supplemental Table 1).

**Fig 2:**
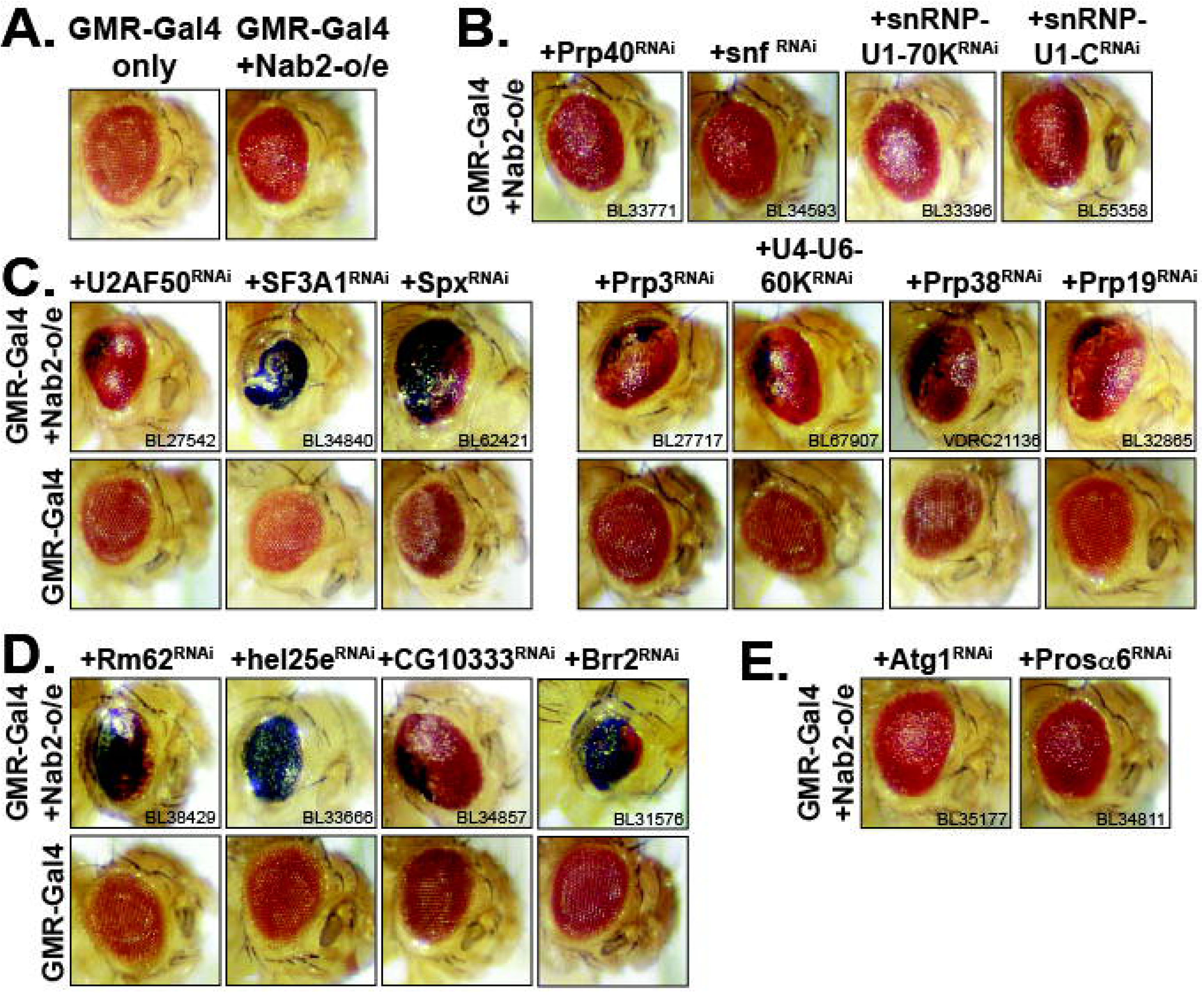
*Nab2* genetically interacts with genes encoding spliceosome components. (A) Overexpression of Nab2 in the fly eye causes eye disorganization and morphology defects (*ninaE.GMR-GAL4/+; Nab2^EP3716^/+*). Control flies (*ninaE.GMR-GAL4/+)* have wild-type eye morphology. Control flies or flies overexpressing Nab2 in fly eyes were crossed to flies harboring UAS-RNA interference (RNAi) constructs targeting (B) components of the U1 snRNP, (C) the U2 snRNP and U4/U5.U6 tri-snRNP, or (D) several RNA helicases implicated in splicing. While reduction of U1 snRNP proteins does not genetically modify Nab2 induced eye morphology defects, decreased expression of U2 and U4/U5.U6 components and selected RNA helicases enhanced eye roughness. Reduced expression of these proteins alone, in the context of wild-type Nab2 levels, caused no or very limited eye morphology defects. (E) Eye-specific reduction of Atg1 and Pros-α6 by RNAi does not dominantly modify the rough-eye phenotype caused by Nab2 overexpression.

Fly eye morphology defects caused by Nab2 overexpression were also enhanced when levels of U4/U6.U5 tri-snRNP components were reduced. Decreased expression of Prp3 (Mount and Salz 2000; Cho et al. 2002) and U4-U6-60K (also sometimes referred to as Prp4 (Mount and Salz 2000)), key components of the tri-snRNP, caused enhanced eye roughness and the presence of posterior eye cell death (as shown by black eye tissue). Reduced levels of Prp31, Prp6, or Prp8, additional components of the U4/U6.U5 tri-snRNP did not modify *Nab2* eye phenotypes (Supplemental Table 1). Reduced expression of Prp38 also caused enhancement of the Nab2-dependent eye roughness phenotype. Prp38 is associated with the U4/U6.U5 tri-snRNP and suggested to be involved in its activation during splicing (Xie et al. 1998; Andersen and Tapon 2008). Importantly, reduced levels of Prp38, Prp3, and U4-U6-60K in fly eyes on their own caused no changes to eye morphology (Fig 2C).

The extensive panel of genetic interactions presented thus far have implicated Nab2 in U2 and U4/U6.U5 snRNP function. During spliceosome activation, U2 and U4/U6.U5 complexes are dramatically reorganized by RNA helicases to create the B^act^ and C complexes (Cordin and Beggs 2013). To test whether *Nab2* also genetically interacts with genes encoding RNA helicases required for spliceosome rearrangement and activation, we reduced levels of selected RNA helicases using RNAi and looked for modification of the fly eye roughness phenotype caused by Nab2 overexpression (Fig 2D). RNAi-mediated reduction of Rm62, Hel25e (also called Sub2 in *S. cerevisiae*, (Eberl et al. 1997; West and Milgrom 2002)), CG10333 (also called Prp28, (Strauss and Guthrie 1994; Mount and Salz 2000; Herold et al. 2009)), and Brr2 (also called l(3)72Ab (Park et al. 2004; Herold et al. 2009)) each caused significant enhancement of eye roughness in flies overexpressing Nab2. Reduced expression of Rm62, Hel25e, and CG10333 caused no phenotype in control eyes containing wild-type Nab2 levels. Reduced expression of Brr2 caused a very subtle alteration in eye morphology of control flies in only one of the RNAi lines used (Supplemental Table 1). Although previous studies have demonstrated that ZC3H14 physically associates with members of the THO complex, a multi-protein complex required for transcriptional elongation (Katahira 2012; Morris and Corbett 2018), reduction in the expression of THO complex members Tho2, cuff, thoc5, or thoc7 did not modify the Nab2 eye roughness phenotype (Supplemental Table 1).

Genetic interaction screens have been utilized widely in *Drosophila,* the budding yeast *Saccharomyces cerevisiae* (where Synthetic Genetic Array (SGA) screens have been widely adopted (Tong et al. 2001; Tong and Boone 2006; Boone et al. 2007)), and now in the form of CRISPR screens in cultured human cells (Fielden et al. 2025; Wu et al. 2025). However, since the spliceosome is such a critical component of gene expression, we wanted to ensure that overexpression of Nab2 in fly eyes was not merely sensitizing eye cells to the loss of any essential gene, such as those involved in autophagy or the ubiquitylation system. Importantly, knockdown of either Atg1, a required component of the autophagy pathway, or Pros-α6, a component of the ubiquitin proteasome, did not dominantly modify the Nab2-overexpression induced rough eye phenotype (Fig 2E and Supplemental Table 1). Finally, although Nab2 does not appear to directly regulate the expression of SmB protein levels when analyzed by immunoblot (Fig1B and Fig 1C), we also analyzed whether Nab2 loss in either larval or adult brains (see below and (Jalloh et al. 2023) for details) was associated with changes in expression of other genes modifying the Nab2 rough eye phenotype using RNA-sequencing data (Supplemental Table 1, (Jalloh et al. 2023), and see below for analysis of RNA-seq data). Fisher’s exact tests indicate that genes modifying Nab2-dependent eye phenotypes were not differentially expressed when Nab2 was lost from larval (p = 0.7223), adult female (p-value = 0.4834), or adult male (p-value = 0.274) brain tissue (Jalloh et al. 2023). The lack of genetic modification with components of the autophagy and ubiquitin pathways and the specificity of the interactions between Nab2 and specific splicing proteins (for example, with SmB/D3/E/F but not with SmD1/D2), strongly suggest that Nab2 may regulate or monitor the quality control of splicing or be more directly involved in spliceosome maturation or recruitment.

### Nab2 loss causes changes to gene expression patterns within developing brain tissue

Previous work has demonstrated that *Drosophila* Nab2 is required for brain development (Kelly et al. 2016; Corgiat et al. 2021; Corgiat et al. 2022; Jalloh et al. 2023), correct dendritic branching patterns (Corgiat et al. 2021), and overall nervous system function (Pak et al. 2011; Bienkowski et al. 2017; Rha et al. 2017). In order to better understand the involvement of Nab2 in splicing and how Nab2 might regulate gene expression and splicing during brain development, we performed whole-brain RNA-seq on RNA samples isolated from late (wandering L3) larval stage brains lacking Nab2 (genotype: *Nab2^ex3^/Nab2^ex3^*) and wild-type control brains (genotype: *Nab2^pex41^/Nab2^pex41^*). Close inspection of aligned reads from these experiments using the Integrative Genome Viewer (IGV, Broad Institute) demonstrated that an extremely small number of reads mapped to the *Nab2* locus in samples from Nab2 null larval brains (Supplemental Fig 1A), confirming the identity of each null sample. The *Nab2^ex3^* allele was derived by imprecise excision of a transposable element originally located within the Nab2 promoter. This imprecise excision resulted in a ∼1.5kb deletion and created the RNA and protein null *Nab2^ex3^* allele (Pak et al. 2011). Overall, each biological replicate generated 26-35 million uniquely mapped reads (Supplemental Data Table 2).

After accounting for batch effects, differential expression (DE) analysis demonstrated that the expression of 1,167 genes was significantly decreased while expression of 1,073 genes was significantly elevated (adjusted p-value < 0.05, Supplemental Data Table 3). Restricting our analysis to those genes that have a log_2_ fold change (log_2_FC) < -1 or >1 revealed that the expression of only 86 genes was significantly elevated, while the expression of 96 genes was decreased in Nab2 null larval brains (Fig 3A). Principal component analysis (PCA) revealed that biological replicates were grouped by genotype along principal component 1 (PC1), suggesting that the primary source of variance could be attributed to genotype (Supplemental Data Fig 1B.) In order to validate the changes in gene expression that were observed by Illumina short-read RNA sequencing, we isolated RNA from an additional 12 biological replicates (6 replicates per genotype, each of which are pools of 60-75 brains) and used the nCounter system (Geiss et al. 2008) to hybridize RNA to a custom designed gene panel containing ∼100 genes that were expected to be increased, decreased, or show no change between Nab2 null and control brain tissue. This system allows for direct quantification of RNA levels in each sample without PCR amplification biases (Geiss et al. 2008). All genes selected for the predicted increased or predicted decreased categories were differentially expressed in our RNA-seq experiment (adjusted p-value < 0.05). As shown in Fig 3B, most genes whose expression was significant decreased or a significant increased when analyzed by RNA-seq were also consistently changed when using the nCounter system (also see Supplemental Table 4). Likewise, the majority of genes (26/31) with very little change in expression between Nab2 null and wild-type control brains in the RNA-seq data remained unchanged in new RNA preparations using the nCounter system (Fig 3B, middle panel). Correlations in gene expression changes could now also be calculated between the RNA-seq and nCounter methods (Fig 3C). A strong correlation was observed when the changes in gene expression between control and Nab2 null samples were compared between these two methods (r = 0.89, gray line in Fig 3C). Using a combination of Illumina short-read RNA sequencing and nCounter experiments, we have demonstrated that Nab2 loss causes a specific set of genes to be consistently over- or under-expressed during central nervous system (CNS) development.

**Fig 3:**
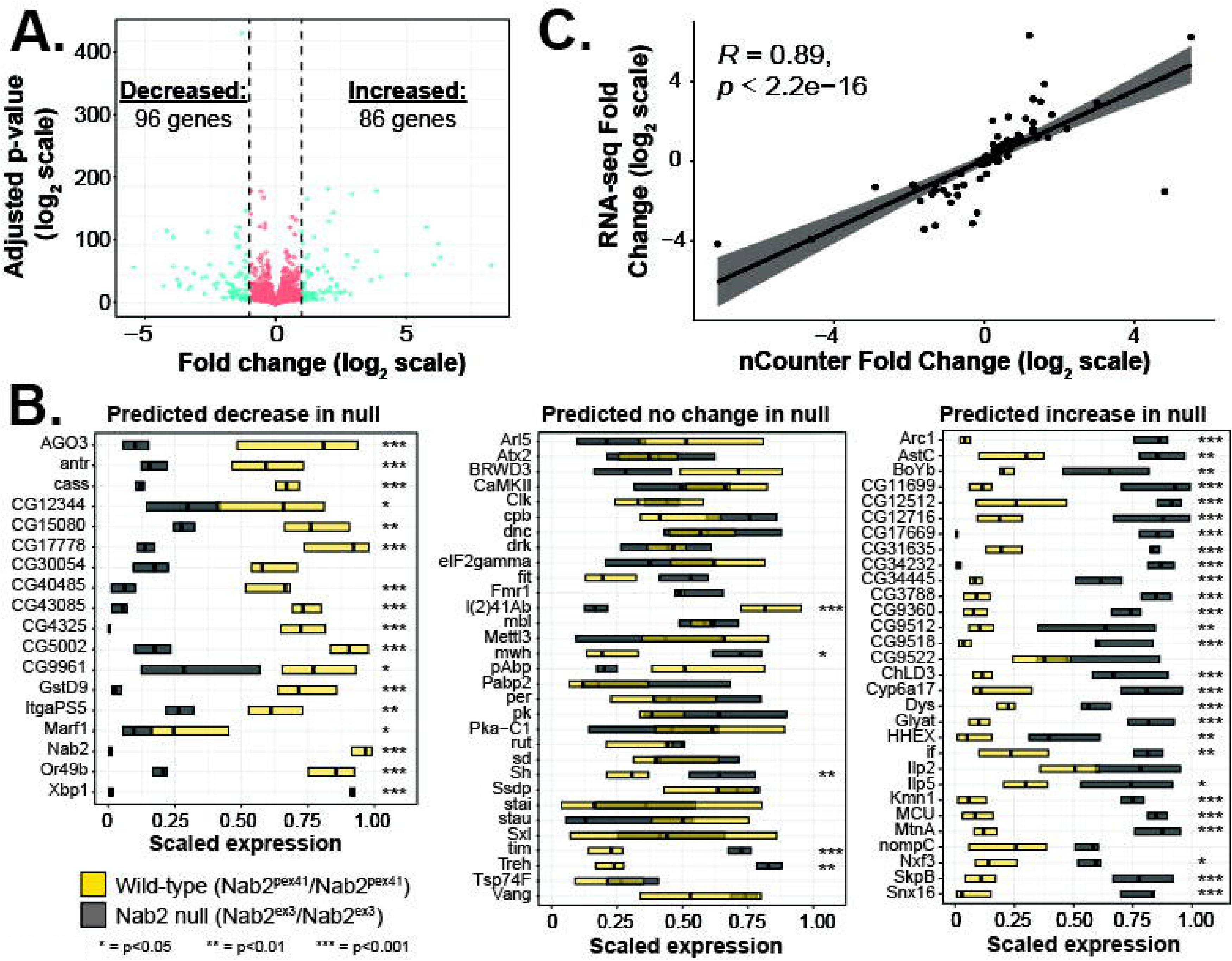
Nab2 regulates gene expression in the developing *Drosophila* brain. (A) Nab2 is responsible for regulating gene expression. Volcano plot shows average fold-change in gene expression between Nab2 null and control L3 larval brain tissue. Genes that were significantly changed more than 2-fold in Nab2 null (*Nab2^ex3^/Nab2^ex3^*, n = 7) and control wild-type (*Nab2^pex41^/Nab2^pex41^*, n = 6) brain tissue compared to control tissue (adjusted p-value < 0.05, log2-fold change > 1 or < -1) are shown in blue. The expression of 96 genes was significantly less abundant in Nab2 null brain tissue, while the expression of 86 genes was significantly elevated. (B) NanoString analysis of gene expression in Nab2 null and control larval brain tissue. Isolated RNA was incubated with a custom designed CodeSet containing genes in one of three categories (predicted decrease, predicted no change, predicted increase) based on RNA sequencing results. Boxplots show the 25^th^-75^th^ percentile and median level of scaled expression for each gene. Asterisks represent genes where significant changes in gene expression were observed by limma using a generalized linear model (* = adj p-value < 0.05, ** = adj. p-value < 0.01, *** = adj. p-value < 0.001). (C) Comparison of RNA-sequencing data and normalized nCounter data revealed a strong level of correlation between fold change values from each experiment (Pearson’s r = 0.89, p < 0.001). Gray shaded areas around the regression line represent the 95% confidence interval.

Using the list of differentially expressed genes generated by DESeq2, we next used clusterProfiler (Subramanian et al. 2005; Yu et al. 2012; Xu et al. 2024) to investigate whether Nab2 loss caused significant changes in the expression of gene sets associated with specific biological processes. Nab2 loss caused changes in the expression of genes (log_2_ fold change >1 or <-1, p<0.05) associated with specific gene ontology (GO) terms such as monooxygenase activity (GO: 0004497), heme binding (GO: 0020037), 17-beta-hydroxysteroid dehydrogenase activity (GO: 0072582), and fatty acid ligase activity (GO: 0015645 and GO: 0120515) (Supplemental Table 5 and Supplemental Fig 2). The lack of enrichment for GO terms associated with a specific biological process may suggest that Nab2 is involved in a fundamental step of RNA biosynthesis that is common to mRNA transcripts encoding proteins from a variety of pathways.

### Nab2 loss causes changes in alternative splicing and intron retention

Given the extensive network of genetic interactions between *Nab2* and components of the *Drosophila* spliceosome (Fig 1 and Fig 2), we next tested the hypothesis that Nab2 loss causes changes in splice site usage within larval brain tissue. Using the ι14′ (Delta Psi, Change in Percent Spliced In) function of the Modeling Alternative Junction Inclusion Quantification (MAJIQ) algorithm (ver. 2.5, (Vaquero-Garcia et al. 2016; Vaquero-Garcia et al. 2023)), we identified and categorized splicing modules that were significantly different between Nab2 null and wild-type larval brain tissue (Fig 4A and Supplemental Table 6). Within significantly changed splicing modules, the most prevalent type of splicing change observed in Nab2 null larval brain samples was intron retention (n = 361), followed by alternative 5’ splice site usage (n = 164) and the use of alternative first exons (n = 149). Additional changes in 3’-splice site usage (n = 75), cassette exons (n = 52) and simultaneous changes in both 5’- and 3’-splice sites (n = 43) were also identified. MAJIQ also identified splicing modules containing different first and last exons and putative alternative 5’- and 3’-splice sites (as described by (Vaquero-Garcia et al. 2023)). Putative first and last exons are created when a splice site is identified that creates a new exon (Vaquero-Garcia et al. 2023).

**Fig 4:**
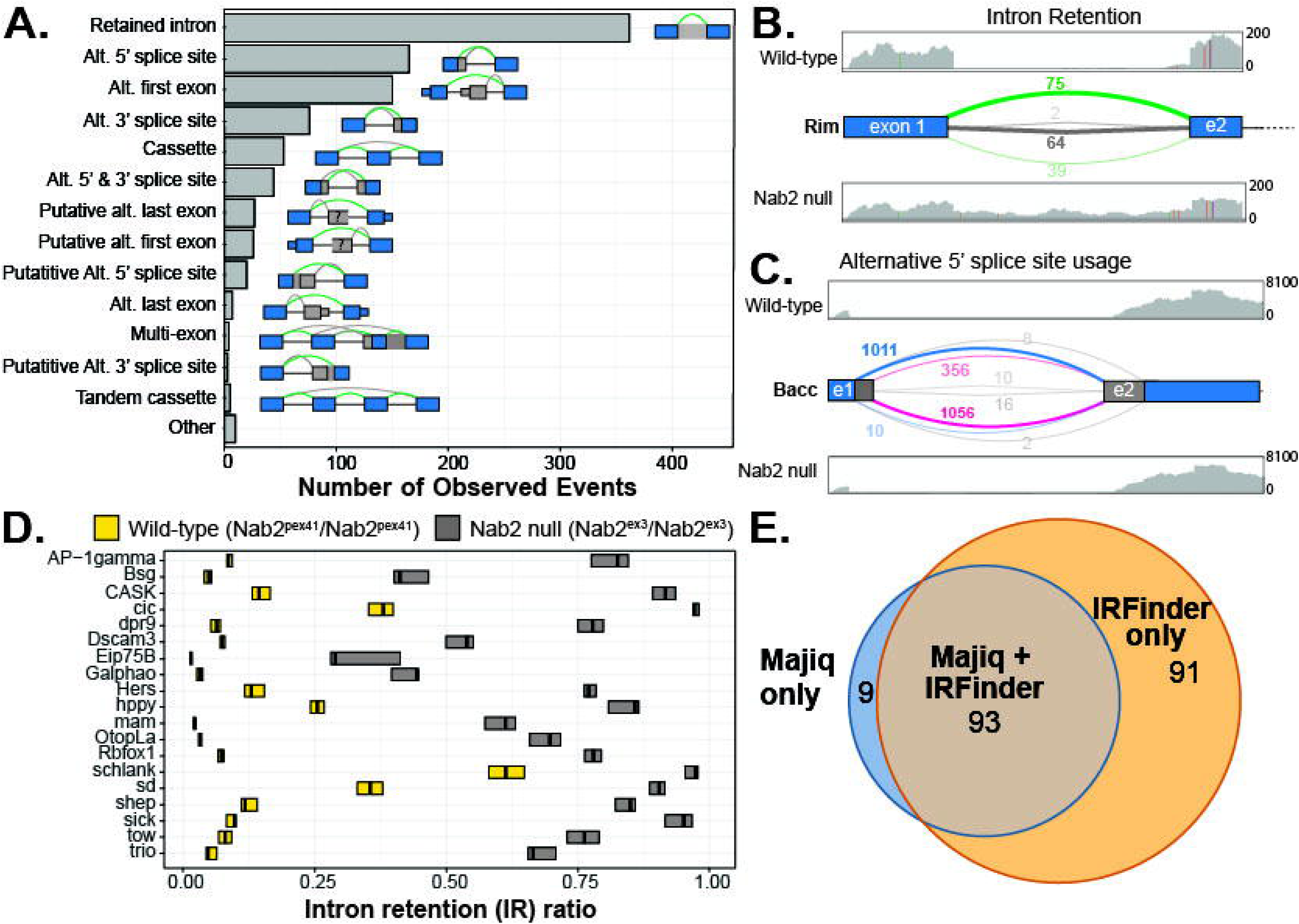
Nab2 loss causes intron retention and changes to alternative splicing patterns. (A) Nab2 is required for the correct regulation of alternative splicing in developing *Drosophila* brain tissue. Aligned RNA-sequencing data from wild-type (*Nab2^pex41^/Nab2^pex41^*) and Nab2 null (*Nab2^ex3^/Nab2^ex3^*) larval brains was used by the Modeling Alternative Junction Inclusion Quantification (MAJIQ) algorithm to quantify the difference in percent selected index (Δψ) between genotypes for each local splicing variation (LSV, (Vaquero-Garcia et al. 2016)). MAJIQ modulizer (Vaquero-Garcia et al. 2023) was then used identify splicing modules were significant differences in splicing had occurred between genotypes. The number of observed splicing events within significantly changed modules is shown. Blue boxes represent annotated exons. Thick gray boxes represent retained introns or changes to exon starting/ending coordinates. Complex splicing events where more than one change in splicing was identified by Majiq could be quantified in multiple categories. (B) MAJIQ splicegraphs and Integrative Genome Viewer (IGV) track screenshots for the fly *Rim* gene demonstrate an increase in first intron retention in Nab2 null brains. Example IGV tracks show sequencing read alignments to specific genomic loci as gray bars or histograms. Exons of *Rim* transcripts are shown as blue boxes, while introns are shown as lines connecting exons. RNA-seq reads from control tissue align to exons but not to introns, while reads from Nab2 null tissue align to both exons and introns. The median number of reads from control samples (median = 75 reads, n = 6 replicates) and Nab2 null samples (median = 39 reads, n = 7 replicates) that support the shown exon 1 – exon 2 splice junction are shown in green near the curved lines of the MAJIQ splicegraphs. The normalized number of reads supporting intron retention from control and Nab2 null tissues are shown in gray/black along the intron. (C) Nab2 is required for the control of alternative splicing of *Bacc* transcripts. IGV tracks reveal read alignment to the *Bacc* locus in either control or Nab2 null samples. 5’ exons of the *Bacc* gene are shown in blue and alternative splicing patterns are represented by the curved lines between exon 1 (e1) and exon 2 (e2). Alternative 5’ and 3’ splice sites, along with extended exons, are shown in gray. Alternative splicing of the e1-e2 splice site causes 3 annotated isoforms to be produced. In wild-type larval brain tissue the upstream 5’ splice site is preferentially used (median of 1011 reads for control samples vs. median of 10 reads per Nab2 null sample). Brains lacking Nab2 preferentially express *Bacc* transcripts that are spliced with the downstream 5’ splice site. A median of 1056 reads per sample support the use of this splice site in Nab2 null larval brains while a median of 356 reads per sample support the use of this splice site in control larval brains. (D) IRFinder validated intron retention events in Nab2 null larval brain tissue. Aligned RNA-seq reads from wild-type control and Nab2 null larval brains were used by IRFinder to calculate intron retention ratios for annotated introns (Middleton et al. 2017). (E) There is a high degree of overlap in the identification of genes expressing transcripts that contain a retained intron in Nab2 null larval brains between Majiq and IRFinder. 92 of the 101 genes identified by Majiq as producing a transcript with a retained intron were also identified as retained by IRFinder.

Inspection of splicegraphs generated by MAJIQ from control larval brain tissue and Nab2 null larval brain tissue shows intron retention in the first intron of the *Rim* gene (Fig 4B). As shown in IGV plots (Fig 4B), there are very few reads that align to the first intron of *Rim* in wild-type brain tissue while RNA-seq reads from Nab2 null larval tissue align with the genomic region between *Rim* exon 1 and exon 2 (e2). The splicegraph generated by MAJIQ (Fig 4B) displays the median number of reads within a group (control or Nab2 null) that cross a given splice junction or that are present within an intron. Although the first intron of *Rim* is correctly spliced in the majority of wild-type samples (*Nab2^pex41^*, a median of 75 reads per sample cross the exon 1 – exon 2 boundary), the first intron is more commonly retained in Nab2 null larval brains. A median of 39 reads cross the *Rim* exon 1 – exon 2 junction in Nab2 null larval brains (*Nab2^ex3^/Nab2^ex3^*), while a normalized median of 64 reads per sample align to intronic sequences in intron 1 of *Rim*. Changes in 5’ and 3’ splice site usage were also observed for other genes upon Nab2 loss. For example, in the first exon of the *Bacc* gene (Fig 4C) there are two annotated adjacent 5’-splice sites. In wild-type larval brains, the 5’-most splice site is preferentially used, while in Nab2 null larval brains, the slightly more downstream 5’-splice site was preferred. A median of 1056 reads among Nab2 null samples support the use of the downstream 5’-splice site while only 356 reads support use of this site in wild-type control larval brain tissue. Thus, Nab2 is implicated in the regulation of multiple types of alternative splicing.

To more comprehensively identify introns that are retained in Nab2 null flies, we next used an additional algorithm, called IRFinder (Middleton et al. 2017), to identify retained introns in our RNA-seq data. Importantly, IRFinder identified many of the same retained introns as MAJIQ in *Nab2^ex3^/Nab2^ex3^*null larval brains (Fig 4D, Fig 4E, and Supplemental Table 6). Overall, 90% (93/102) of the genes identified by Majiq as producing a transcript with a retained intron were supported by IRFinder (Fig 4E and Supplemental Data Table 6). In summary, we have used two independent algorithms to identify a consistent set of transcripts with retained introns in Nab2 null brains, suggesting that Nab2 is necessary for accurate splice site selection and the prevention of intron retention.

To validate the bioinformatics approaches used to identify retained introns caused by Nab2 loss, we next used PCR to amplify cDNAs generated from wild-type larval brain RNA samples or RNA samples derived from Nab2 null larval brains. PCR primers (shown as red lines above the gray boxes in Fig 5A) were designed to anneal to exon sequences on either side of the retained intron and therefore would produce different length products depending on whether splicing had occurred. In wild-type larval brains, very few reads aligned to the shown intronic regions while reads are clearly aligned with annotated exons (as shown in IGV plots, Fig 5A). Similarly, the most prominent PCR product in wild-type samples corresponded to the shorter, spliced product (denoted with an “S”) present near the bottom of each gel (Fig 5A). Except for *Trio* isoform D and Ca-α1D, less than 10% of transcripts from the genes analyzed here contain the intronic region in wild-type controls (Fig 5C). Several transcripts, such as the A and C isoforms of the *Trio* gene (Fig 5A), existed as a collection of alternatively spliced transcripts and were therefore represented by multiple, slightly less prominent bands (denoted as “Alt Spl” in Fig 5A). Nab2 loss primarily caused an increase in the number of RNA-seq reads aligning to intronic regions, as shown by the gray bars in IGV screenshots (Fig 5A, top). Lack of Nab2 also caused an increase in the amount of PCR product produced from the longer unspliced template (denoted with “US” next to each gel) for *Trio* isoforms A and D*, Cask-α, Ca-α1D, Rbfox1, Rim, AP-1γ, and Gα_o_*. Splicing changes were also specific to the affected isoforms and not generalized across all isoforms from the same gene. For example, the longer *CASK-β* and *Gα_o_-B* isoforms, which start from a different transcriptional start site than *CASK-α* and *Gα_o_-A* isoforms, respectively, were not mis-spliced in larval brains lacking Nab2 (Fig 5B). Thus, Nab2 loss from developing brain tissue caused increased intron retention within specific mRNA transcripts.

**Fig 5:**
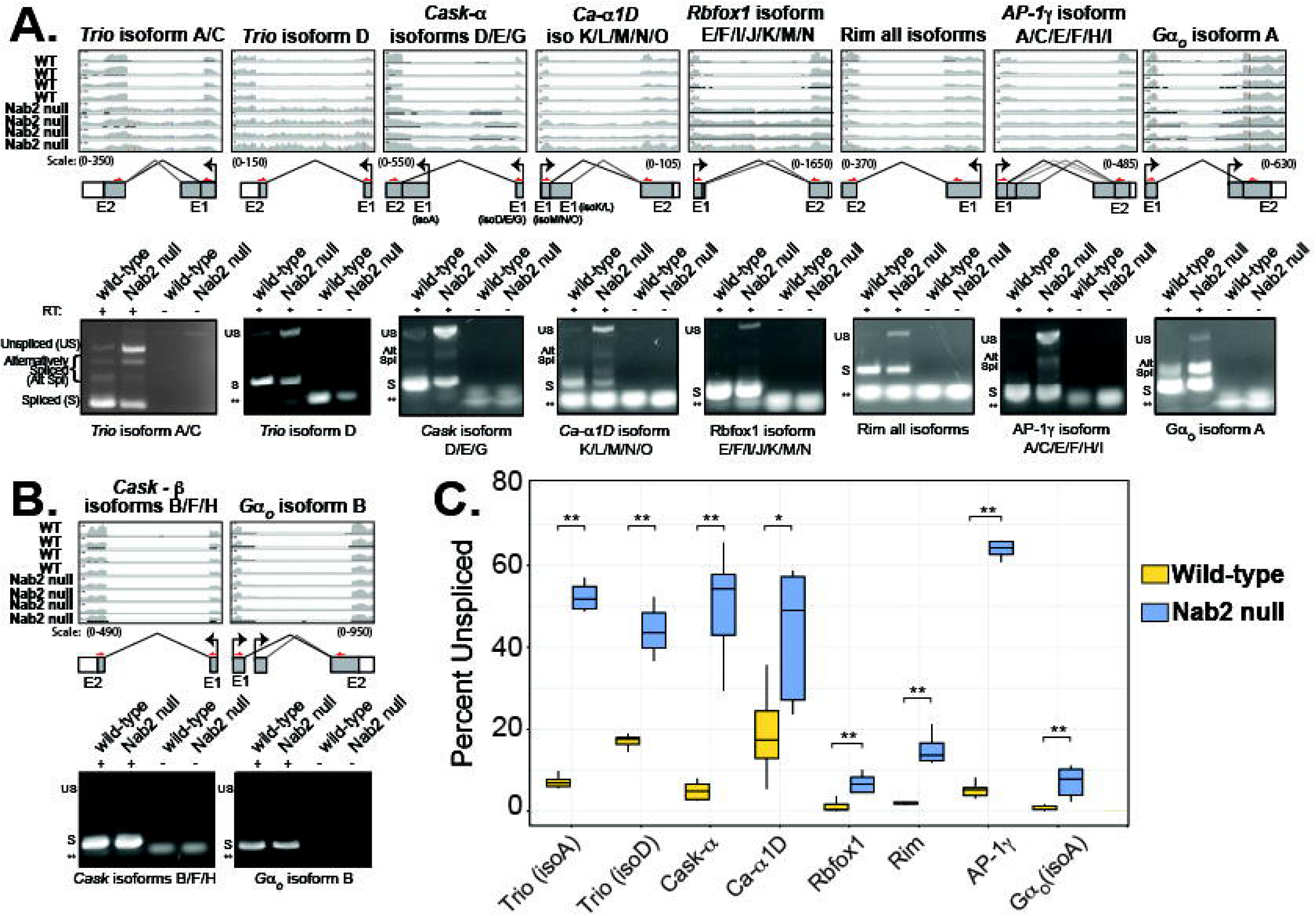
Nab2 prevents intron retention and is required for the regulation of alternative splicing. (A) Semiquantitative reverse transcriptase PCR validates intron retention in Nab2 null larval brains. IGV tracks are shown representing RNA-seq read alignments for four wild-type (WT, *Nab2^pex41^/Nab2^pex41^*) and four Nab2 null (*Nab2^ex3^/Nab2^ex3^*) biological replicates. Exons for each gene are shown below IGV tracks. All IGV tracks for the same isoform were manually scaled to the same values, shown as scale values below each IGV screenshot. Black 90° bent arrows represent transcription start sites for each transcript, gray boxes represent non-coding exons, and white boxes represent coding sequences. Exon 1 (E1) and exon 2 (E2) are labeled. Lines between boxes represent annotated splicing events. Small red arrows represent binding sites for forward and reverse primers used in PCR reactions. RNA from control and Nab2 null larval brains was reverse transcribed with reverse transcriptase (RT+) into cDNA and cDNA was used with gene specific primers to amplify sequences around retained introns. Reverse transcriptase reactions were also performed in the absence of RT enzyme (RT -). PCR products were then resolved on 1-2% agarose gels. Spliced (S), unspliced (US), and alternatively spliced (alt. spl.) PCR products are shown. For some reactions, primer dimers are marked by the double asterisk (**). For each of the shown transcript isoforms, longer PCR products are observed in Nab2 null samples, representing unspliced transcripts. Changes in alternative splicing (Alt. Spl.) are also observed for *Trio*, *Cask-α*, *Ca-α1D*, *AP-1γ*, and *Gα_O_* transcripts. (B) Changes in alternative splicing observed in Nab2 null brains are isoform specific. Semi-quantitative RT-PCR using primers for *CASK-β* and *Gα_O_* isoform B transcripts do not shown evidence of intron retention or changes in alternative splicing in Nab2 null larval brains. (C) Quantification of PCR product signal intensity for spliced vs. unspliced transcripts. The ratio of PCR product intensity corresponding to unspliced transcripts was divided by the intensity of PCR products that correspond to fully spliced transcripts for both control (n = 6) and Nab2 null (n = 7) biological replicates and compared by Wilcoxon tests (* = p<0.05 and ** = p<0.01, using Benjamini-Hochberg corrections to account for multiple comparisons between genotypes).

Changes in mRNA processing can often trigger quality control checkpoints, such as nonsense mediated decay (NMD), that degrade faulty transcripts (Kurosaki and Maquat 2016). Therefore, we reasoned that transcripts containing retained introns in Nab2 null flies might be decreased in abundance. Surprisingly, intron retention caused by Nab2 loss was correlated with increased gene expression (Supplemental Fig 3A). Genes producing a transcript with a retained intron were more likely to be significantly expressed in Nab2 null brain tissue than genes than those genes that did not produce an intron-containing transcript (Fisher’s exact test, p < 0.001). Since changes in mRNA processing caused by Nab2 loss appear to be isoform specific and it can be difficult to discern isoform-level changes in expression from short-read RNA-seq experiments, we next used quantitative real-time PCR with isoform specific primers to assess whether Nab2 loss causes alterations in the levels of select mis-processed isoforms. The expression of *Trio-isoform D* and multiple *Rbfox1* isoforms were significantly elevated in Nab2 null larval brains (Supplemental Fig 3B). Although somewhat elevated, the expression of CASK-α, Ca-α1D, Rim, and *AP-1γ* was not significantly altered.

### Nab2 loss preferentially causes retention of longer first introns

Next, we hypothesized that introns retained in Nab2 null brains might have a common set of characteristics. A preliminary visual inspection in IGV of the retained introns identified by both Majiq and IRFinder initially suggested that Nab2 loss might cause the retention of first introns. Indeed, when the genomic coordinates of the retained introns were used to identify where in the transcript each intron was located, 91/95 retained introns were the first intron of their transcripts (Fig 6A). Since the length of an intron may help determine how 5’- and 3’-splice sites are recognized by the spliceosome (Fox-Walsh et al. 2005; Li et al. 2019; Carranza et al. 2022), we next analyzed the length of retained introns. Although most fly introns, regardless of position within a transcript, were approximately 100 nucleotides in length (median length of all introns was 104 nucleotides (nts), n = 152,810 introns), the median length of all first introns from all transcripts was significantly longer (median = 341 nts, n = 27,957 first introns, p < 0.001, green in Fig 6B). Contrary to other studies where retained introns were on average shorter than constitutive spliced introns (Green et al. 2020; Porras-Tobias et al. 2025; Faretra et al. 2026), the length of retained introns in Nab2 null brains was significantly longer than first introns or all introns (median = 834 nts, n = 91 introns, p <0.001). In addition, first exon length was also significantly shorter (median = 202 nts) for transcripts with retained first introns than a typical first exon (median = 234 nts, p < 0.05, Fig 6C). Since splice site strength is one predictor of intron retention (Monteuuis et al. 2019), we compared the strength of all 5’- and 3’-splice sites of retained introns to all *Drosophila* first introns using MaxEntScan (Yeo and Burge 2004). As demonstrated in Fig 6D, both the 5’ and 3’ splice sites of retained introns were significantly weaker than the average splice site within the *Drosophila* genome. Thus, Nab2 prevents the retention of longer introns with significantly weaker splice sites within developing fly brain tissue.

**Fig 6:**
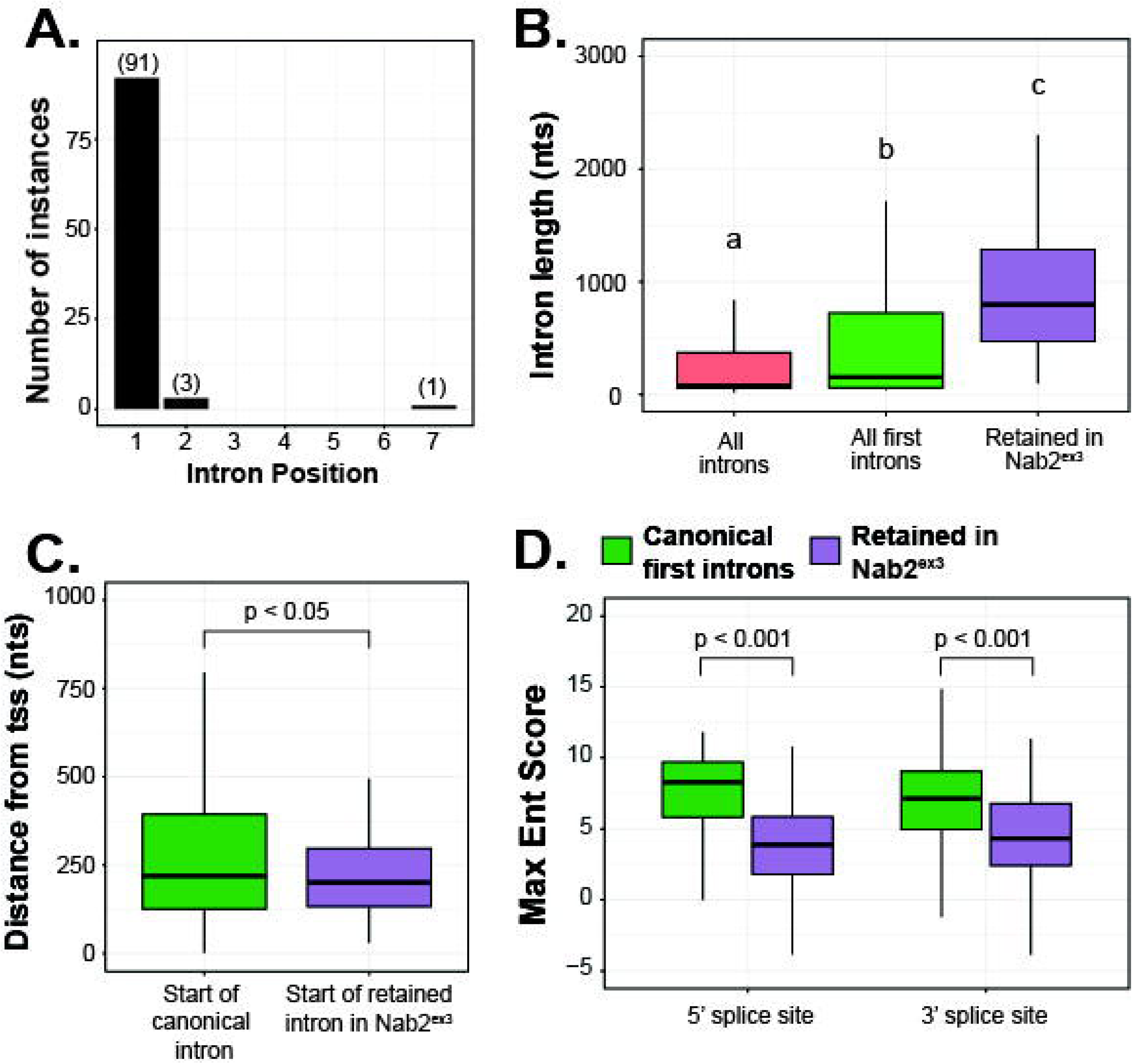
Nab2 prevents retention of long first introns with weak splice sites. (A) Nab2 loss caused retention of first introns. A total of 95 retained introns were identified by both Majiq and IRFinder by coordinate overlap. Several genes express more than one isoform with a retained intron. The number of times an intron was found at a specific position within a transcript is shown in parentheses. (B) Introns retained in Nab2 null larval brains (median length = 834 nts, n = 91 introns) were significantly longer than an average *Drosophila* intron (median length = 104 nts, n = 152,810 introns) or an average first intron in *Drosophila* (median length = 341 nts, n = 27,957 introns). All groups were statistically different from one another, as determined by pairwise Wilcoxon tests; µ was set to the median of the “all introns” dataset (p < 0.001, represented by letters a-c above each boxplot). (C) The median length of first exons from all annotated coding transcripts (median length = 234 nts, n = 27,957 exons) was significantly longer than the length of exons upstream of retained first introns in Nab2 null larval brains (median = 202 nts, n = 91) according to a Wilcoxon test (p <0.001, µ = mean length of all first exons). Data is plotted as the distance from the transcription start site (tss) to the end of the first exon. (D) Nab2 loss causes retention of introns with weaker splice sites. Splice site scores were obtained using a Maximum Entropy (MaxEnt) model, MaxEntScan (Yeo and Burge 2004). Lower scores signify a lower probability that a given sequence functions as a splice site. 5’-splice sites for retained introns in Nab2 null larval brains (median score: 3.77, n = 91) had significantly lower MaxEntScan scores when compared by a one sample Wilcoxon test to the splice sites for all first introns (median score: 8.23, n = 27,957, p < 0.001, µ = median 5’ splice site score for all first introns). Likewise, 3’-splice sites for retained introns in Nab2 null larval brains (median score: 4.24, n = 91) also had significantly lower MaxEntScan scores when compared by a Wilcoxon test to the splice sites for all first introns (median score: 7.11, n = 27,957, p < 0.001, µ = median 3’ splice site score for all first introns).

### 3’ splice sites from retained introns are rich in adenosine

Since Nab2 and its human ortholog, ZC3H14, bind preferentially to poly(A) sequences (Kelly et al. 2007; Pak et al. 2011), we reasoned that Nab2 might associate with A-rich sequences near splice sites to influence splicing efficiency. If true, sequences adjacent to the 5’- and 3’-splice sites of introns retained in Nab2 null brains might be enriched for adenosine. We compared the adenosine content of the 150 nucleotides (nts) surrounding the 5’ and 3’ splice sites of retained introns to that of all first introns in the *Drosophila* transcriptome (Fig 7A). There was a significant increase in adenosine content in exon sequences upstream of the 5’ splice site of retained introns compared to all first intron 5’ splice sites (Fig 7B, p<0.05). There was no significant difference in the adenosine content downstream of the 5’ splice site, within the retained first intron. A rolling average of adenosine content near 5’ splice sites also demonstrated a slight increase in the percentage of As between retained and all introns in the exon upstream of the 5’ splice site (Fig 7C). The 5’ splice site of most (90/91) retained first introns contained the canonical GU dinucleotide but most lack a strong consensus sequence in the immediately upstream exon (Figure 7D, arrowhead marks the exon/intron boundary).

**Fig 7:**
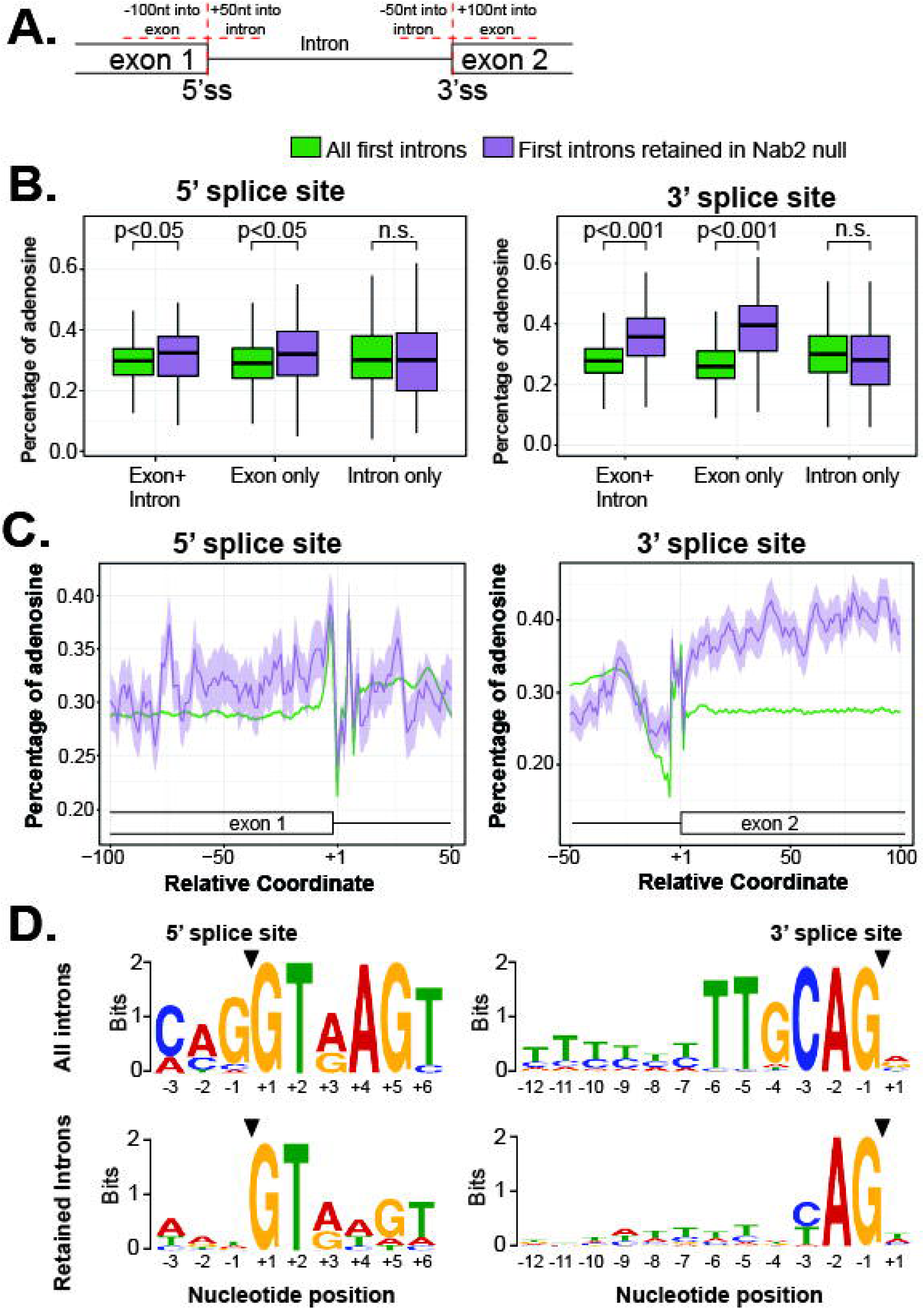
Exons downstream of retained introns contain significantly more adenosine than a typical second exon. (A) Diagram depicting regions of analysis. (B) Exons near retained first introns have significantly higher adenosine content than average *Drosophila* first introns. For 5’ splice sites (left), the percentage of adenosine in exon sequences (100 nts upstream of the 5’ss), intron sequences (50 nts downstream of 5’ss), and exon + intron sequences was quantified for all *Drosophila* first introns (green) and first introns retained in Nab2 null larval brain tissue (purple). For 3’ splice sites (right), the percentage of adenosine in exon sequences (100 nts downstream of 3’ss), intron sequences (50 nts upstream of 3’ss), and exon + intron sequences was quantified. More adenosine was present in exons surrounding introns that were retained in Nab2 null brain tissue, but not in the retained introns themselves (median percent of adenosine for each region is shown, p-values derived from Wilcoxon tests). (C) Five nucleotide rolling average of adenosine content near 5’ (left side) and 3’-ss (right side) of introns retained in Nab2 null tissue demonstrates more adenosine in exon regions, especially in the 100 nts downstream of 3’ splice sites. (D) Retained introns in Nab2 null brains have required 5’ (GT or GU) and 3’ (AG) splice sites but are less likely to have typical upstream and downstream sequences. The 9 DNA nucleotides around 5’-splice sites (-6 to +3, where +1 is first nucleotide of the intron) and the 23 nucleotides surrounding the 3’-splice sites (-20 to +3, where -1 = last nucleotide of the intron) were used for MEME analysis of over-represented sequences. Sequence logos show the probability of each letter at each position in the sequence, scored in bits. Larger letters represent a higher likelihood that the letter will be found at that position relative to the frequency of all possible letters at that position.

Sequences surrounding the 3’ splice sites of retained first introns also had significantly more adenosine than typical first intron 3’ splice sites (Fig 7B, p<0.001). A sliding window analysis (Fig 7C) demonstrated that the percentage of adenosine within the sequence upstream of the 3’ss was similar between retained and all introns until immediately before the 3’ splice site. Here, the adenosine content decreased in all first introns near the poly-pyrimidine tract but remained higher in the retained introns (Fig 7C). Consistent with this finding, pyrimidine-rich sequences (cytosine and thymine/uracil) were not as prevalent in retained introns between positions -11 and -5 immediately before the 3’ splice site (Fig 7D, arrowhead marks the exon/intron boundary). Interestingly, exon 2 sequences downstream of retained first introns (Fig 7C, right side, purple line) contained a significantly higher percentage of adenosine than typical second exons (Fig 7C, green line). The percentage of adenosine in the first 100 nts of exon 2 downstream of a retained intron was nearly 40%, while the adenosine percentage of a typical second exon was approximately 27% (Fig 7B, p < 0.001). Together, these data suggest that introns retained in Nab2 null brains are significantly longer than average, contain significantly weaker 5’- and 3’-splice sites, and lack a strong poly-pyrimidine tract adjacent to the 3’ splice site. Furthermore, exons downstream of retained first introns contain significantly more adenosine than a typical *Drosophila* second exon.

### Introns retained in Nab2 null developing fly brains are enriched in genes required for morphogenesis, axon guidance, and synapse function

If Nab2-regulated intron retention is used to coordinate the expression of several genes required for the same biological process, Nab2-regulated genes might be associated with gene ontology (GO) terms related to that process. Using clusterProfiler (Yu et al. 2012), we found that genes producing transcripts with retained introns in Nab2 null developing fly brain tissue (Supplemental Fig 4A, 4B, and Supplemental Table 5) were significantly enriched for gene ontology terms related to morphogenesis (GO: 0007560 and GO: 0007552), regulation of neurotransmitter secretion and transport (GO: 0046928 and GO: 51588), axon guidance (GO: 0007411), cell adhesion (GO: 0007155), and synapse assembly (GO: 0007416). Several genes, such as *Trio*, *CASK*, and GαO appear as connection points between related biological processes required for nervous system development and function (Supplemental Fig 4B). Thus, our GO term analysis supports a model where Nab2-dependent regulation of intron retention might serve as a mechanism for coordination of related biological processes that are involved in nervous system development.

### Long read RNA-sequencing suggests that Nab2 loss from larval brain tissue causes first intron retention in otherwise fully processed mRNAs

Our data thus far has demonstrated that Nab2 loss causes a substantial number of alternative splicing changes. Until recently, however, Nab2 has been primarily implicated in the control of poly(A) tail length. Thus, we next wanted to investigate whether Nab2 might coordinate these two processing events on the same mRNA transcripts. To supplement our short-read Illumina sequencing, we next utilized Oxford Nanopore Technologies (ONT) sequencing to investigate both intron retention and poly(A) tail length in RNA samples from Nab2 null and wild-type control larval brains. RNA samples were isolated from freshly dissected Nab2 null and control larval brains and either sequenced directly (using direct RNA (dRNA) sequencing) or used to synthesize cDNA, which was then sequenced (cDNA sequencing). Importantly, cDNA synthesis for ONT sequencing was performed using anchored oligo-d(T) primers that captured poly(A) tail length for all transcripts.

Although ONT sequencing of cDNA generated substantially more reads than dRNA sequencing, the average quality score, read length, and percentage of reads aligned to the *Drosophila* genome were comparable between each type of sequencing (Supplemental Table 2). Visual inspection of the *Nab2* locus using IGV demonstrates a clear absence of aligned reads near the transcription start site of the *Nab2* gene (Supplemental Fig 5), consistent with our short-read RNA sequencing (Supplemental Fig 1). A small number of reads that align with the 3’- end of the *Nab2* locus appear to originate from an upstream non-coding RNA.

Visual inspection of dRNA ONT sequencing reads from Nab2 null brains confirmed retained introns in many of the loci identified by IRFinder and Majiq. For example, *AP-1γ* and *Rbfox1* were both identified as having retained first introns in Nab2 null brains (Fig 5D). ONT sequencing demonstrated that while most transcripts produced from the *AP-1γ* and *Rbfox1* genes in wild-type brains are correctly spliced, many transcripts from these genes in Nab2 null brains retain the first intron (Supplemental Figure 5). Interestingly, sequencing reads in both control and Nab2 null samples support annotated splice site junctions in more downstream regions of *AP-1γ* and *Rbfox1* transcripts, demonstrating that transcripts containing a retained first intron in Nab2 null larval brain tissue are otherwise correctly spliced.

### Genes producing transcripts with retained first introns also produce transcripts with extended poly(A) tails in Nab2 null larval brains

In addition to providing information about isoform-specific splicing, both cDNA and dRNA ONT sequencing can provide an estimate of poly(A) tail lengths for each sequenced transcript. We utilized both types of ONT sequencing in our poly(A) tail length analyses here to ensure that estimates derived after cDNA synthesis matched those observed in the direct RNA sequencing protocol. The median transcriptome-wide poly(A) tail length detected by ONT sequencing was similar between wild-type and Nab2 null samples and between the methods of sequencing (cDNA: 76 nts in wild-type, 78 nts in Nab2 null; dRNA: 79 nts in wild-type, 76 nts in Nab2 null).

Although bulk tail length was similar between wild-type and Nab2 null samples, we reasoned that Nab2 loss might preferentially regulate the poly(A) tail length of transcripts containing a retained first intron. Unfortunately, the 3’ bias observed in our ONT sequencing datasets, the 5’ location of the retained first introns observed in Nab2 null brain tissue, and the average length of our sequencing reads (Supplemental Table 2) prevented us from reliably analyzing tail length and intron retention on the same reads. Therefore, all poly(A) tail length estimates were derived from reads aligned at the gene level, not by isoform. Using this analysis and looking first in wild-type flies, the median poly(A) tail length was 76 nts for transcripts whose splicing was not regulated by Nab2 (Fig 8A, Nab2-independent genes). Genes, such as *Rbfox1*, *Trio*, and *AP-1γ*, that produce a transcript where first intron retention is regulated by Nab2 have a significantly longer median poly(A) tail length of 104 nts (p < 0.001). Interestingly, the median poly(A) tail length of transcripts produced from most genes does not change when Nab2 is lost (median poly(A) tail length = 78 nts). However, Nab2-regulated genes produce transcripts with significantly longer poly(A) tails in Nab2 null tissue (median poly(A) tail length = 124 nts, p < 0.001). Almost identical results are observed in a single replicate of dRNA sequencing (Fig 8B). Similar to our cDNA sequencing results, dRNA sequencing demonstrated that genes with an intron whose splicing was regulated by Nab2 contained longer poly(A) tails (median tail length of 79 nts vs. 100nts in wild-type tissue, p<0.01). The tails of these transcripts became even longer in Nab2 null samples, when Nab2 was missing and intron retention occurred (median tail length of 76nts vs. 113 nts, p<0.01). Using our cDNA sequencing data, we next analyzed median poly(A) tail length changes for each of the genes that produce an intron-retained mRNA in Nab2 null brains. As shown in Figure 8C, 65% of Nab2 regulated genes produce transcripts with poly(A) tails that increased by more than 10 nts (Fig 8C), while tail length remains relatively unchanged (<10 nts difference, shown in gray) for transcripts from ∼30% of Nab2-regulated genes. Upon Nab2 loss, only 5/88 Nab2-regulated genes produce transcripts with shorter tails. This data suggests that genes producing mRNAs where intron retention is regulated by Nab2 also produce transcripts that have extended poly(A) tails upon Nab2 loss.

**Fig 8:**
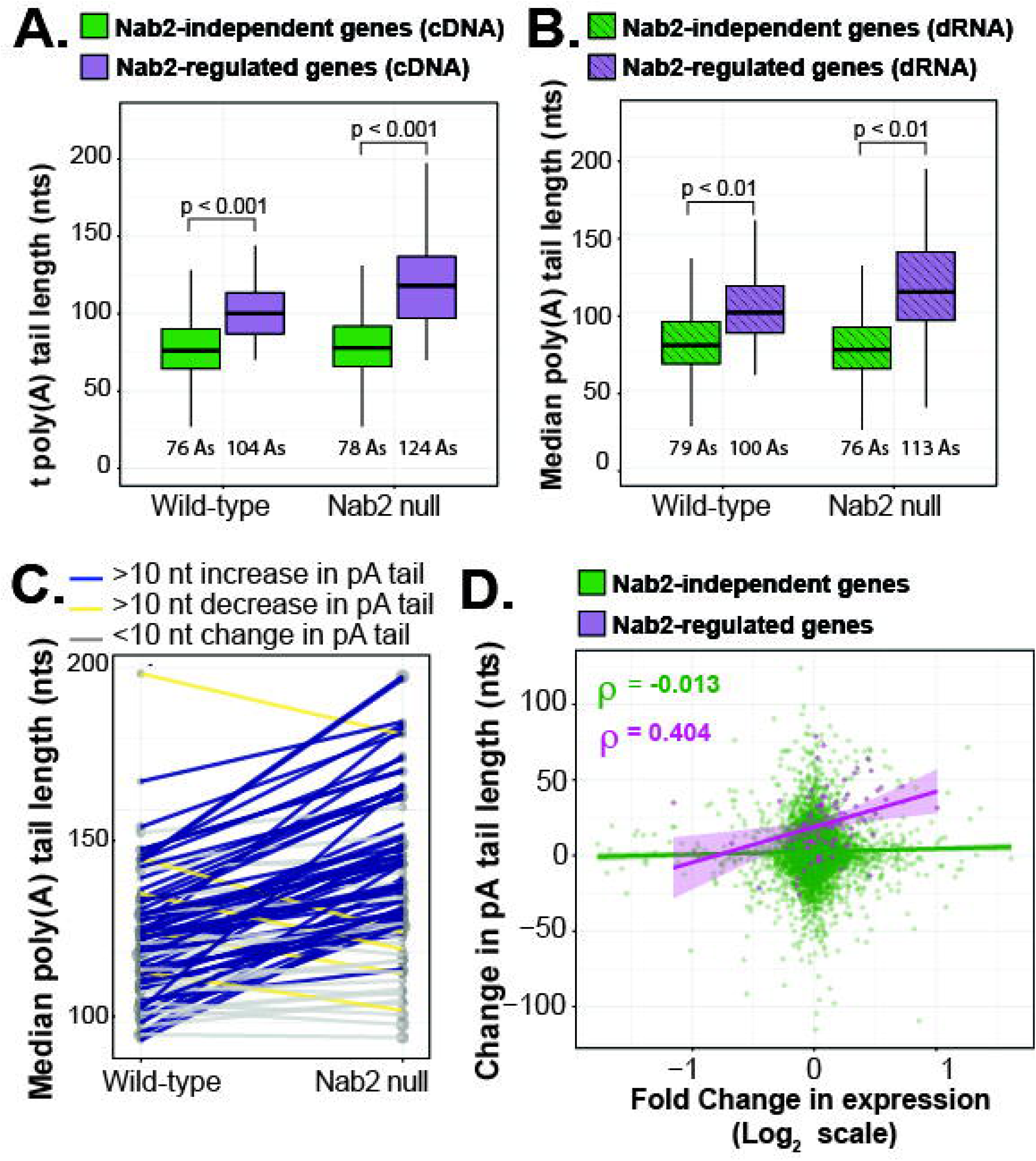
Genes producing transcripts containing retained first introns in Nab2 null larval brains also produce transcripts with longer poly(A) tails. RNA was collected from freshly dissected control/wild-type (*Nab2^pex41^/Nab2^pex41^*) and Nab2 null (*Nab2^ex3^/Nab2^ex3^*) larval brains and sequenced by (A) Oxford Nanopore cDNA sequencing or (B) direct RNA (dRNA) sequencing. Genes were separated into two groups: Nab2-regulated genes that produce transcripts containing a retained first intron in Nab2 null larval brains (purple) and Nab2-independent genes, which do not have any intron-retention events in Nab2 null brains (green). The median poly(A) tail length for all genes in each category is shown underneath the boxplots. (A) For cDNA-ONT sequencing, the median poly(A) tail length of Nab2-regulated genes (n = 88 genes, median tail length = 104 nts) is significantly longer than Nab2-independent genes (n = 8,317 genes, median tail length = 76 nts) in wild-type brains (p < 0.001, Wilcoxon test, W = 591500). In Nab2 null brain tissue, there is a significant increase in the poly(A) tail length of transcripts from Nab2-regulated genes (n = 88 genes, median tail length = 124 nts) compared to genes that do not produce intron-containing transcripts in Nab2 null brains (n = 8,154 genes, median tail length = 78 nts, p < 0.001, Wilcoxon test, W = 637988). (B) Similarly, when direct RNA (dRNA) sequencing was used, Nab2-regulated genes have significantly longer poly(A) tails in wild-type tissue (Nab2-independent genes: n = 6,870 genes, median tail length = 79 nts; Nab2-dependent genes: n = 88 genes, median tail length = 100 nts, p <0.001, Wilcoxon Test, W = 449084). Poly(A) tail lengths are even longer for Nab2-regulated genes in larval brains lacking Nab2 (Nab2-independent: n = 6,841 genes, median tail length = 76 nts; Nab2-dependent: n = 86 genes, median tail length = 113 nts, p < 0.001 by Wilcoxon test, W = 490096). (C) Median poly(A) tail lengths for all Nab2-regulated genes are shown in ONT cDNA sequencing samples from wild-type and Nab2 null brains. Genes producing transcripts with longer poly(A) tails (> 10nts longer in Nab2 null) are shown in blue (n = 57 genes), genes producing transcripts with poly(A) tails that are at least 10 nts shorter in Nab2 null brains are shown in yellow (n = 5 genes), and genes that have similar length tails (less than 10 nts difference) between wild-type and Nab2 null samples are shown in gray (n = 26 genes). (D) The fold-change in gene expression between wild-type and Nab2 null brain tissue was plotted as a function of poly(A) tail length difference between wild-type and Nab2 null samples. A positive correlation between increased gene expression and increased poly(A) tail length in Nab2 null larval brains was observed for Nab2-regulated genes (purple, n = 91, rho = 0.404), but not for Nab2 independent genes (green, n = 6,630, rho = -0.013).

Thus far, we have shown that Nab2 appears to regulate a subset of mRNA transcripts. Nab2 loss causes retention of first introns and extended poly(A) tails on these transcripts. We hypothesized that changes in gene expression observed in Nab2 null larval brains (as determined from DeSEQ2 analysis of short-read RNA Seq data) may be correlated with changes in poly(A) tail length (as determined from long-read cDNA ONT sequencing data). Importantly, we investigated this question for all genes (Fig 8D, green) and then for genes that produce a Nab2 regulated transcript (Fig 8D, purple). No correlation (rho = -0.013) between gene expression changes and poly(A) tail differences in Nab2 null flies was observed for all transcripts. However, when we narrowed our focus to only those genes where intron retention is regulated by Nab2, a moderate positive correlation (rho = 0.404) was observed between changes in gene expression and poly(A) tail length, suggesting that for transcripts containing retained introns, longer poly(A) tails in Nab2 null brains are correlated with larger changes in gene expression. Collectively, this suggests that Nab2 may coordinate multiple mRNA processing events, such as intron retention and poly(A) tail length control, with overall expression of target mRNA transcripts.

### Intron retention causes significant alterations in encoded protein levels and localization within the developing *Drosophila* brain

To test whether increased intron retention observed in Nab2 null brains would cause changes in protein abundance, soluble proteins were isolated from dissected control and Nab2 null larval brain tissue and the levels of three candidate proteins, Trio, Rbfox1, and Dystroglycan (Dg), were analyzed. The retained first introns in both *Trio* and *Rbfox1* transcripts are located in the 5’-untranslated region (5’ UTR) of the transcript. *Dg* expresses four alternatively spliced isoforms in *Drosophila* and Nab2 loss from developing fly brain tissue is predicted to change the typical pattern of Dg isoform expression.

Trio functions as a guanine exchange factor (GEF) for Rac GTPases during brain development (Awasaki et al. 2000; Newsome et al. 2000; van Rijssel and van Buul 2012) and is highly expressed within the fly mushroom body, a brain structure required for olfactory memory formation (Awasaki et al. 2000; Lin 2023). Since the retained *Trio* introns are located in the 5’ UTR of *Trio* transcripts, we wondered whether intron retention caused by Nab2 loss would significantly alter Trio protein levels. Although loss of Nab2 causes intron retention in *Trio* RNA isoforms A/C and D (Fig 6A), only isoform D protein levels were significantly decreased in Nab2 null brains (Fig 9A, top panel). Loss of Nab2 caused a 46% decrease in the levels of Trio- isoform D (p < 0.05), but no significant change in the levels of Trio-isoformA/C protein (Fig 9B) was observed (p > 0.05). To assess whether loss of Nab2 caused localized changes in Trio expression, we dissected control and Nab2 null larval brains and used immunofluorescence to visualize Trio protein location and abundance. GFP was also expressed using the GAL4/UAS system (Brand and Perrimon 1993) to highlight mushroom body neurons. At the larval stage, Trio was expressed throughout the developing brain but was enriched in mushroom body neurons (Fig 9C and (Awasaki et al. 2000)). However, in brains lacking Nab2, Trio was no longer concentrated in mushroom body neurons but can still be visualized as diffuse staining signal in other areas of the brain.

**Fig 9:**
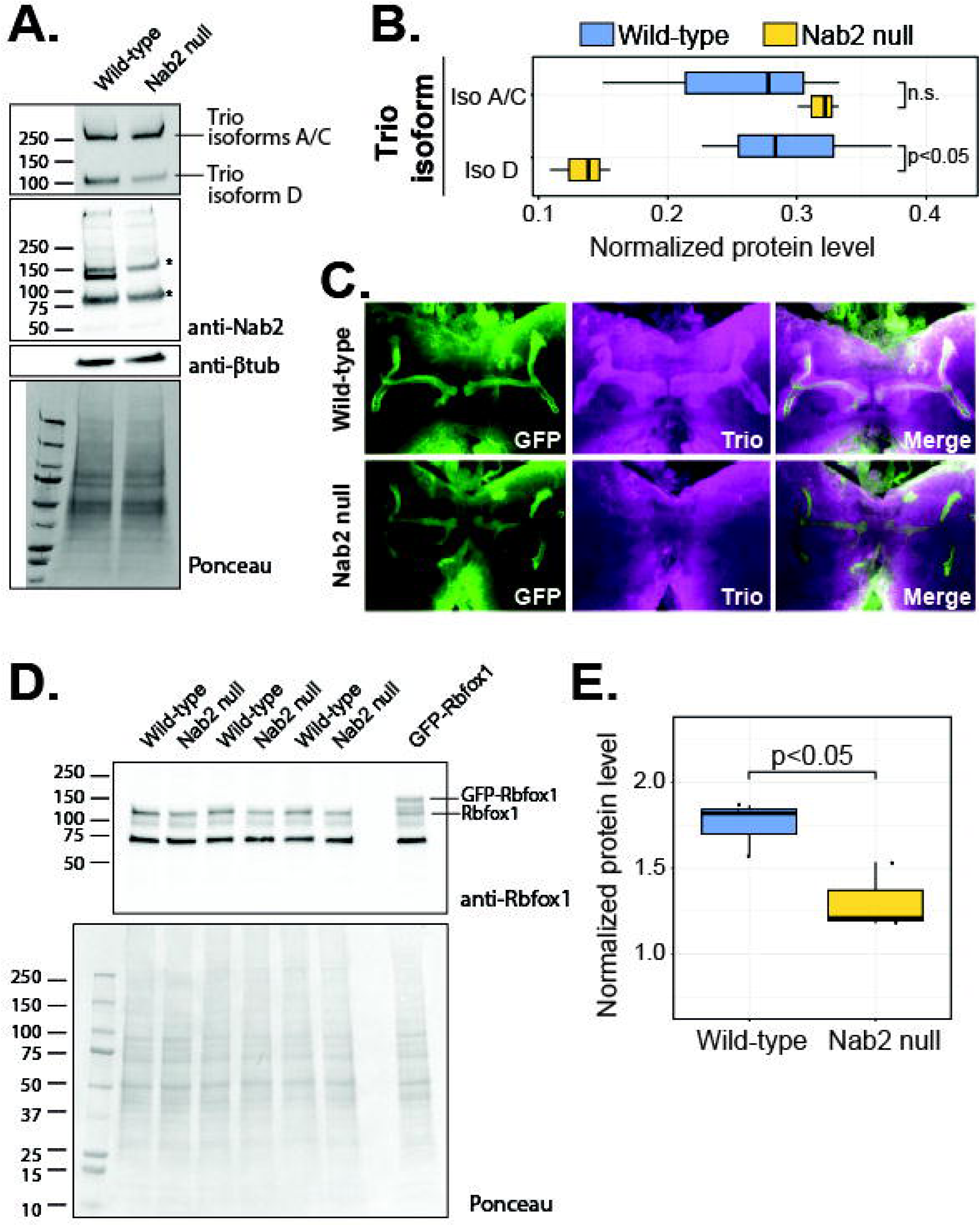
Trio and Rbfox1 protein levels and localization are controlled by Nab2 during larval brain development. (A) Nab2 is necessary for correct expression of Trio protein isoform D but not Trio isoforms A and C in larval brains. Wild-type control (*Nab2^pex41^/Nab2^pex41^*) and Nab2 null (*Nab2^ex3^/Nab2^ex3^*) brains were dissected from wandering L3 larvae and pooled into batches of 15-17 brains. Soluble protein from lysed brain tissue was then used for immunoblots to detect the level of Trio, Nab2, and β-tubulin protein. Monoclonal Trio antibodies recognized isoforms A and C, which are close in size and indistinguishable here, as well as isoform D. The expected size of Nab2 is approximately 140 kDa. Polyclonal Nab2 antibodies recognized several non-specific bands (marked by *) in addition to Nab2, which is missing in Nab2 null brain lysates. Tubulin was included as a loading control. (B) Quantification of Trio bands in (A), normalized to Ponceau staining. Normalized levels of Trio-isoforms A/C are not significantly different between wild-type and Nab2 null brains, but Trio-isoform D protein levels are significantly decreased in Nab2 null brains (paired t-test, t(4) = -3.5716, p=0.0233, n = 3 for each genotype). (C) Nab2 is necessary for expression of Trio in larval mushroom body brain tissue. Wild-type (genotype: *c305a-GAL4/UAS-CD8-GFP*) and Nab2 null (genotype: *c305a-GAL4/UAS-CD8-GFP;Nab2^ex3^/Nab2^ex3^*) brains expressing GFP in mushroom body neurons were dissected from L3 larvae and Trio protein expression was visualized by immunofluorescence and confocal microscopy. Trio expression is enriched in wild-type mushroom body neurons, but is substantially decreased in larval brains lacking Nab2. (D) Nab2 regulates expression of Rbfox1 protein in larval brain tissue. 15-20 control or Nab2 null larval brains were dissected and pooled, tissue was lysed, and the level of Rbfox1 protein was detected by immunoblot. Three separate biological replicates of 15-20 brains per replicate are shown for each genotype. One pooled sample of ∼15 larval brains expressing GFP-Rbfox1 from the endogenous Rbfox1 promoter is also shown. (E) Quantification of (D), showing normalized levels of the ∼125 kDa isoform of Rbfox1 protein. Rbfox1 was significantly less abundant in Nab2 null brains compared to controls (two-sample t-test, t(4)=3.1066, p = 0.036, n = 3 for each genotype). Smaller molecular weight bands were not quantified. The band at approximately 75 kDa is not predicted to be Rbfox1 (see (Guven-Ozkan et al. 2016), figure S4).

Rbfox1 (also referred to as A2BP1) is an RNA binding protein that has previously been implicated in controlling alternative splicing and intron retention (Lovci et al. 2013; Fu and Ares 2014; Misra et al. 2015) by binding to UGCAUG sequences near splice sites (Jin et al. 2003; Auweter et al. 2006; Ray et al. 2013). Similar to *Trio*, several *Rbfox1* transcripts contain retained 5’UTR introns in Nab2 null brain tissue (Fig 5A). Wild-type and Nab2 null larval brains were dissected, pooled into groups of ∼15 brains, and soluble protein was isolated and used for immunoblotting with a polyclonal Rbfox1 antibody (Tastan et al. 2010; Carreira-Rosario et al. 2016). A sample from flies containing a MiMIC insertion (Venken et al. 2011) expressing GFP-Rbfox1 from the endogenous promoter was also included as a positive control for antibody specificity. GFP-Rbfox1 appeared as a slightly larger band and several smaller isoforms when probed with the Rbfox1 antibody (Fig 9D). A significant decrease in the largest ∼125 kDa isoform of Rbfox1 was consistently observed in Nab2 null larval brain samples (Fig 9E, p < 0.05). The band located at ∼75 kDa was previously predicted to be an unrelated protein often detected by this antibody but unrelated to Rbfox1 (see Fig S4 from (Guven-Ozkan et al. 2016)). Overall, loss of Nab2 during brain development causes intron retention and decreases in Trio and Rbfox1 protein abundance.

### Retained introns contain binding motifs for other RBPs

Since decreases in Rbfox1 levels may contribute to the intron retention observed in Nab2 null brains, we used the Analysis of Motif Enrichment (AME) and the Find Individual Motif Occurrences (FIMO) algorithms (McLeay and Bailey 2010; Grant et al. 2011) to analyze whether Rbfox1 binding sequences were enriched in the retained introns as well as the upstream and downstream 50 nucleotides. Although FIMO found 14 Rbfox1 (also called A2BP1) binding sequences in the retained introns (Supplemental Fig 6A, orange label), Rbfox1 binding sequences were not significantly enriched in the entire data set of retained introns (Supplemental Fig 6B) compared to a background group of all first introns. However, binding motifs for other RBPs, including Bruno1 (also called Arrest/ARET), Smooth (Sm), Papi, and Sex-lethal (Sxl), among others, were significantly enriched in retained intron sequences (Supplemental Fig 6B). Interestingly, the binding site for the cytoplasmic poly(A) binding protein (PABP, binding sequence: GAAAAHV, where H = uracil, and V = G, C, or A nucleotides) was significantly enriched in retained introns (Supplemental Fig 6A), consistent with our data showing an enrichment of adenosine near retained introns (Fig 7). Further experimentation will be necessary to characterize the complement of RNA binding proteins associated with retained introns and also to understand how that group of RBPs might collaboratively work with Nab2 to regulate Nab2-dependent transcripts.

### Nab2 loss causes changes in alternative exon usage

In addition to intron retention, Nab2 loss also caused changes to alternative splicing patterns and isoform switching in select transcripts. For example, differential splicing of *Dg* exons 8 and 9 gives rise to four *Dg* isoforms (Fig 10A and (Schneider and Baumgartner 2008)): isoform A lacks exon 8 (predicted size = 111 kDa), isoform B lacks both exons 8 and 9 (predicted size = 102 kDa), isoform C lacks the smaller exon 9 (predicted size = 130 kDa), and isoform D contains all exons (predicted size = 138 kDa). Splicegraphs generated by MAJIQ (Vaquero-Garcia et al. 2016; Vaquero-Garcia et al. 2023) from our Illumina RNA-seq data suggest that both wild-type and Nab2 null brain tissue express roughly equal amounts of *Dg*-isoform B and very little *Dg*-isoform A (Fig 10B). In wild-type larval brains, a median of 204 reads and 263 reads per sample were detected by MAJIQ that support expression of *Dg*-isoforms C and D, respectively. However, a large increase in the median number of reads supporting *Dg*-isoform C (404 reads compared to 68 reads for Dg-iso D) was observed in Nab2 null larval brains, suggesting that *Dg*-isoform D was not as highly expressed in cells lacking Nab2. To investigate these changes in transcript levels, we performed semi-quantitative PCR using primers that amplify this differentially spliced region (primers anneal in exons 8 and 11, red arrows in Supplemental Fig 7). In strong agreement with our RNA-seq data, wild-type larval brain tissue expressed both *Dystroglycan* isoform C and D, while larval brain tissue lacking Nab2 expressed primarily the shorter C isoform of *Dystroglycan* that lacks exon 9 (Supplemental Data Fig 7B).

**Fig 10:**
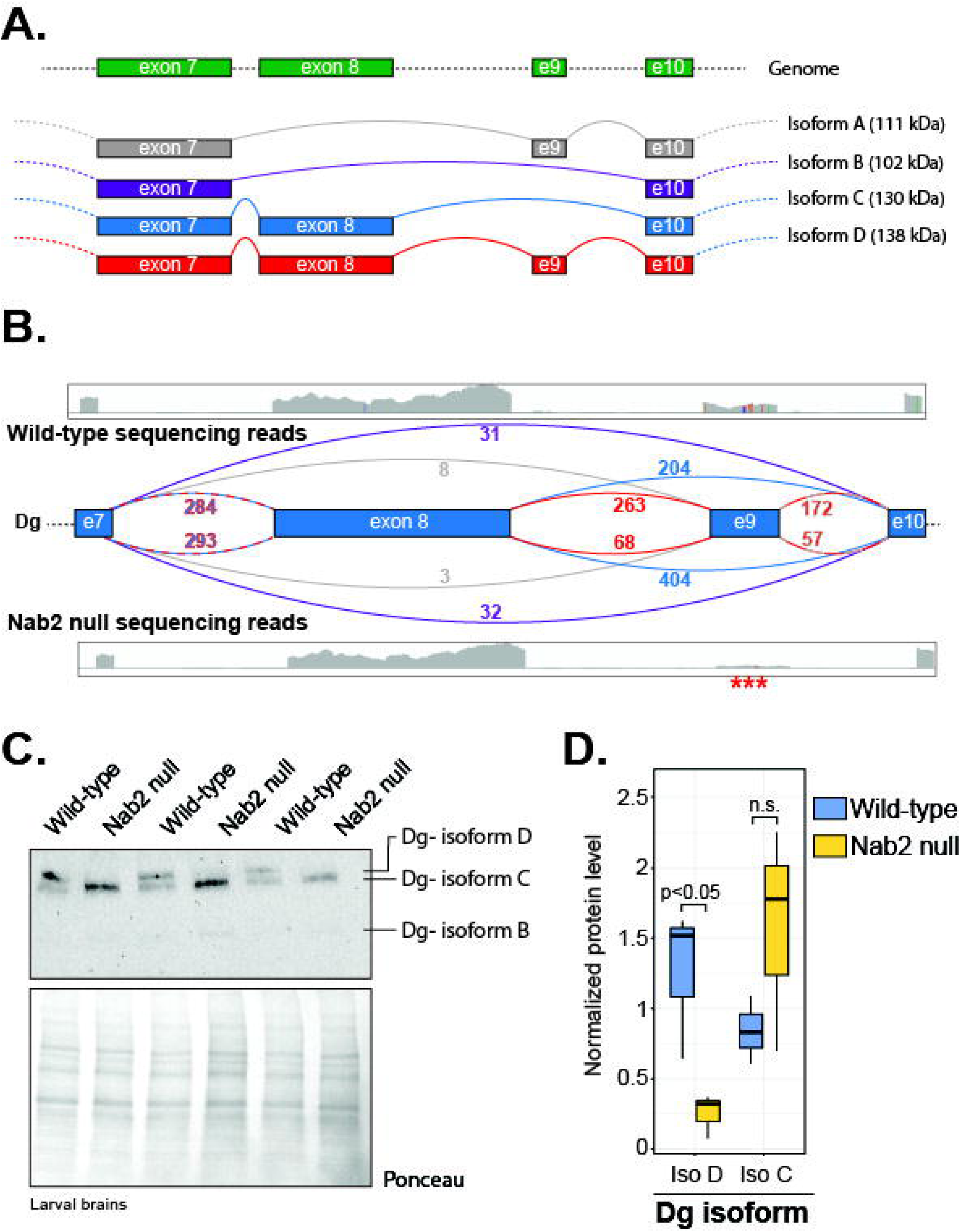
Nab2 is necessary for correct alternative splicing of Dystroglycan isoforms. (A) *Dystroglycan* (*Dg*) encodes four isoforms. The genomic organization of *Dg* exons 7-10 is shown at the top in green (e9 = exon 9, e10 = exon 10) and alternatively spliced transcripts are shown below. Isoform A lacks exon 8 and is predicted to have a molecular mass of 111 kilodaltons (kDa). Isoform B lacks both exon 8 and exon 9 and is predicted to have a molecular mass of 102 kDa. Isoform C lacks exon 9 and is predicted to have a molecular mass of 130 kDA. Isoform D contains all exons and is predicted to have a molecular mass of 138 kDa. (B) IGV screenshots demonstrate the density of aligned reads from wild-type (*Nab2^pex41^/Nab2^pex41^*) or Nab2 null (*Nab2^ex3^/Nab2^ex3^*) larval brain RNA samples to the *Dg* locus. *Dg* exon locations are shown as blue boxes and curved lines connecting exons represent potential splice junctions. The color of the curved line corresponds to the isoform shown in A where that splice junction is used. The junction between exon 7 (e7) and exon 8 is found in *Dg*-isoforms C and D and is therefore dashed in red and blue. The splice junction between exons 9 and 10 is shared between *Dg-*isoforms B and D and is dashed in red and gray. The median number of reads (as determined by MAJIQ) from wild-type larval brain RNA that support each splice junction is shown above the splice junction diagram while the median number of reads from Nab2 null larval brains is shown next to the curved lines below the splice junctions. The number of reads supporting intron retention in each sample has been omitted for clarity. (C) Nab2 regulates expression of Dystroglycan (Dg) protein isoforms. 15-20 control or Nab2 null larval brains were dissected and pooled, tissue was lysed, and the level of Dg protein was detected by immunoblot. Isoforms are labeled based on predicted molecular weights. (D) Quantification of (C), showing normalized levels of Dg-isoforms C and D. Due to low expression levels, Dg isoform B was not quantified. Dg isoform D was significantly less abundant in Nab2 null brains compared to controls (paired t-test, t(4)=-3.1162, p = 0.0356, n = 3 for each genotype). Dg isoform C was also slightly, but not significantly, elevated in Nab2 null brain tissue (paired t-test, t(4)=1.531, p = 0.200, n = 3 for each genotype).

In order to test whether changes in alternative *Dg* splicing patterns caused by Nab2 loss affected Dg protein abundances, we analyzed levels of Dg protein in developing wild-type and Nab2 null larval brain tissue using an antibody that recognizes all Dg isoforms (Deng et al. 2003). Both control and Nab2 null brain tissue expressed very low levels of Dg-isoform B protein (Fig 10C). Dg-isoform A was not expressed in developing brains at detectable levels. Control brain tissue expressed roughly equal amounts of Dg-isoform C and Dg-isoform D, while brain tissue lacking Nab2 preferentially expressed only the shorter C isoform of Dg (Fig 10C, D). In summary, Nab2 regulates alternative splicing of cassette exons within *Dg* mRNA transcripts. Nab2 loss caused significant changed in Dg protein isoform abundance within developing brain tissue.

## Discussion

In the current study we have demonstrated that the *Drosophila melanogaster* RNA binding protein Nab2, an evolutionarily conserved ortholog of human ZC3H14, functions as a critical regulator of alternative splicing and polyadenylation during brain development. Nab2 genetically interacts with multiple components of the spliceosome, and Nab2 loss causes significant changes in gene expression, alternative splicing, and 5’-UTR intron retention during brain development. Nanopore sequencing suggests that transcripts containing retained introns are otherwise fully spliced and that Nab2 loss may cause extension of poly(A) tails on intron-containing transcripts. Finally, changes in alternative splicing of *Trio*, *Rbfox1*, and *Dg* transcripts caused by Nab2 loss result in alterations in protein abundance and isoform usage, which might collectively result in the pleiotropic effects previously observed in Nab2 null flies, including alterations in mushroom body development (Kelly et al. 2016) and viability defects (Pak et al. 2011 2014). Together, these data highlight a new role for Nab2 in the regulation of intron retention during brain development and suggest that Nab2 may coordinate mRNA splicing and polyadenylation of specific target transcripts.

The genetic interactions between *Nab2* and components of the spliceosome suggest that Nab2 may function in either snRNP maturation or at a point in RNA processing that might affect all splicing steps. First, *Nab2* functionally interacts with genes encoding Sm proteins, core components of all snRNPs and critically required proteins in spliceosome function (Matera and Wang 2014). Sm proteins initially assemble into smaller, well-ordered sub-complexes before coming together into a holo-Sm ring structure (Neuenkirchen et al. 2015). These complexes include a dimer of SmD1 and SmD2, a dimer of SmB and SmD3, and a trimer of SmE, SmF, and snRNPG (SmG) (Raker et al. 1996; Fischer et al. 2011). *Nab2* genetically interacts with genes encoding components of the SmB/D3 and Sm E/F/G complexes, but not with genes encoding the SmD1/D2 complex (Fig 1). In addition, Nab2 phenotypes are not modified by knockdown of pICln or Smn (Supplemental Table 1 and Fig 1), chaperones required for the ordered assembly of snRNP complexes. Together, the observed genetic interactions suggest that Nab2 might regulate snRNP maturation or function, but it seems unlikely that Nab2 loss completely disrupts formation of the snRNPs required for mRNA splicing.

Given that Nab2 overexpression in the fly eye is enhanced by reduction in the expression of genes that produce proteins required for multiple splicing steps (Supplemental Table 1), it seems unlikely that Nab2 only regulates a single step in the splicing process. For example, reducing the expression of Sf3a and Sf3b members might impact early steps in A complex formation and U2 snRNP recruitment, while reducing the expression of Prp3 or U4-U6-60K could lead to destabilization of or failure to recruit the U4/U6.U5 tri-snRNP complex. Therefore, rather than regulating a single step in splicing and given the propensity of Nab2 and ZC3H14 to bind poly(A) RNA (Kelly et al. 2007; Pak et al. 2011), several non-mutually exclusive models could explain the observed changes in alternative splicing in Nab2 null larval brains.

First, Nab2 might physically contact A-rich sequences within exons downstream of weak splice sites and recruit the spliceosome to promote efficient splicing. Once processed, exon junction complex (EJC) proteins could be deposited and the spliced mRNAs exported to the cytoplasm for translation. Loss of Nab2 would prevent efficient recruitment of the spliceosome to weaker adenosine-rich splice sites and subsequently lead to intron retention. Although we have not yet tested whether *Drosophila* Nab2 physically associates with spliceosome proteins, yeast Nab2 binds the splicing factor Mud2, suggesting that conserved physical associations between Nab2 and spliceosome components may exist. Second, our data also demonstrates that Nab2 regulates the abundance of several RNA binding proteins and splicing factors, including Rbfox1 (Fig 9), U4-U6-60K, and Hel25e (Supplemental Table 3). Previously work has established the role of Rbfox1 in alternative splicing (Lovci et al. 2013; Mukherjee and Nongthomba 2023) and reduction of U4-U6-60K (also called Prp4) causes intron retention in *timeless* (*tim*) transcripts in *Drosophila* (Shakhmantsir et al. 2018). Thus, changes in the abundance of these proteins might also indirectly contribute to intron retention that occurs in Nab2 null tissue. Additional experiments will be required to comprehensively assess whether changes in the abundance of U4-U6-60K, Hel25e, or Rbfox1 have an additive or synergistic effect on the splicing alterations observed in Nab2 null larval brains.

A final model suggests that Nab2 might coordinate RNA modifications and RNA processing. Nab2 loss was previously shown to elevate levels of N^6^-methyladenosine (m6A) at specific sites within the *Drosophila* transcriptome (Jalloh et al. 2023; Lancaster et al. 2024). Changes in m6A abundance have been linked to the regulation of alternative splicing (Louloupi et al. 2018; Zhou et al. 2019; Jiang et al. 2021; Zhu et al. 2023; Santos-Rodriguez et al. 2026) and in flies, m6A is most abundant in the 5’-UTR regions of mRNA transcripts (Kan et al. 2021). Surprisingly, the vast majority of retained introns in Nab2 null brains are first introns and most are located in 5’-UTRs. Thus, Nab2 loss might increase m6A deposition within the 5’-UTR of a target transcript and subsequently prevent or alter recruitment of the spliceosome, leading to increased intron retention or changes in exon inclusion in the final mRNA. A preliminary analysis of our data suggests that approximately one-third of the retained introns in Nab2 null larval brains contain an annotated m6A site (using m6A data from adult heads (Kan et al. 2021)), supporting this model. However, the effects of m6A deposition can often vary by transcript. Usually, m6A is deposited by a complex containing the “writer” protein, Mttl3, and is then physically bound by a “reader” protein, such as the nuclear Ythdc1 protein (Jiang et al. 2021). In some cases, binding of Ythdc1 then recruits the splicing regulator SRSF3 (Xiao et al. 2016; Jiang et al. 2021) and helps to increase the likelihood of exon inclusion into the mature mRNA. However, other studies have demonstrated that the 3’ splice site AG sequence can be methylated by a different methyltransferase, Mettl16, to prevent recognition by the spliceosome (Mendel et al. 2021). In addition, studies in *Drosophila* have shown that m6A controls splicing of *Sxl* transcripts to regulate production of Sxl protein (Granadino et al. 1990; Hilfiker et al. 1995; Kan et al. 2017; Jalloh et al. 2023), which is a main determinant of female sex determination in flies (Bell et al. 1988; Duan et al. 2026). In females, deposition of m6A prevents inclusion of a male-specific exon into *sxl* transcripts and promotes the production of full-length Sxl protein (Duan et al. 2026). Future studies investigating interactions between Nab2, specific RNA sequences near retained introns, and components of the spliceosome and m6A “writers” will be necessary to develop a more comprehensive understanding of how Nab2 regulates the retention of first introns within the 5’-UTR regions of target transcripts.

Intron retention is a prevalent mechanism of alternative splicing that has the potential to significantly alter the levels of an encoded protein. However, when retained introns are present in 5’- or 3’-UTRs it is difficult to predict how a retained intron will impact protein levels. In our current study, intron retention within specific mRNAs, such as *Trio*-isoforms A/C or specific *Rbfox1* isoforms, occurs within the 5’-UTR and decreases the abundances of the encoded proteins. While changes to the non-coding regions of a transcript might be expected to have little or no effect on the abundance of the encoded protein, previous studies have demonstrated that 5’-UTR alterations can significantly impact ribosome recruitment to reporter RNAs (Lewis et al. 2025; Plassmeyer et al. 2025) and may lead to changes in protein production. In addition, several neurodevelopmental disorders have been linked to 5’-UTR mutations (Whiffin et al. 2020; Wright et al. 2021), further supporting the importance of 5’-UTR regions in the regulation of translation. Consistent with a previous study showing that extended 5’-UTRs decreased ribosome recruitment to reporter transcripts (Lewis et al. 2025), intron retention, and therefore elongation of 5’-UTR sequences, was associated with a decrease in Trio-A/C and Rbfox1 protein abundance in our study. Although 5’-UTR extension might prolong ribosome scanning and slow translation initiation, it is more likely that 5’-UTR extension also increases the likelihood that inhibitory elements, such as upstream open reading frames (uORFs), microRNA binding sequences, or RBP binding sequences, would be present in the mRNA transcript. Indeed, our data suggests that Nab2-regulated introns are enriched for the binding sequences of several RNA binding proteins (Supplemental Fig. 6), including Aret/Bru1, Sm, Papi, and Sxl, among others. Although Arrest (Aret, also called Bruno1) is well-known to interact with 3’-UTR sequences and repress translation of the *oskar* mRNA during early fly development (Kim-Ha et al. 1995; Webster et al. 1997), it can also regulate splicing events in flight muscle development (Nikonova et al. 2024), suggesting that these RBPs might play a variety of roles in the post-transcriptional regulation of gene expression. In addition, sex lethal (sxl) can also function as a translational inhibitor by preventing translation initiation of the *msl-2* mRNA (Gebauer et al. 2003), suggesting that similar mechanisms might occur on transcripts with retained introns in Nab2 null flies. Future studies focused on identifying the cohort of RBPs bound to retained intron sequences and the functional consequences of RBP binding will help to shed more light on the exact mechanisms by which extended 5’-UTRs and intron retention can regulate protein production.

Previous studies have implicated ZC3H14 and Nab2 in the global, transcriptome-wide control of polyadenylation (Hector et al. 2002; Pak et al. 2011; Grenier St-Sauveur et al. 2013; Kelly et al. 2014; Soucek et al. 2016; Rha et al. 2017; Gabs et al. 2025). Many of the alterations in poly(A) tail length caused by ZC3H14/Nab2 loss are also tissue specific. For example, loss of ZC3H14 expression causes extended poly(A) tails in mouse hippocampus, but not in cortex or liver tissue (Rha et al. 2017). One interpretation of these data is that Nab2 might regulate the abundance and poly(A) tail length of specific transcripts in particular tissues or developmental time points. Our data supports this model by demonstrating that Nab2 loss causes retention of first introns in specific transcripts, that those transcripts are likely to have longer poly(A) tails, and that genes producing these transcripts are more highly expressed in Nab2 null larval brains. Thus, our data extends the previous literature by narrowing Nab2 targets for poly(A) tail length control to those containing retained first introns. Further studies using reporters and a targeted sequencing approach will be necessary to quantify both the abundance and average poly(A) tail length of specific isoforms of Nab2-regulated genes in wild-type and Nab2 null tissue.

Our data showing extensive genetic interactions with spliceosome components and pervasive intron retention upon Nab2 loss suggest that Nab2 could be added to a growing list of RNA binding proteins that coordinate expression of genes required for brain development. In support of this idea, genes encoding transcripts with retained introns are enriched for GO terms associated with synaptic assembly, neurotransmitter secretion, and axon guidance (Supplemental Fig 4). Nab2 also appears to regulate alternative splicing and prevent intron retention within multiple transcripts required for mushroom body development (Lin 2023), including *Trio* (Awasaki et al. 2000), *sickie* (*sick*, (Abe et al. 2014)), *still life* (*sif*, (Ng and Luo 2004)), *Dg* (Yatsenko et al. 2021), *Abrupt* (Kucherenko et al. 2012), and *IGF-II* mRNA binding protein (*Imp*, (Liu et al. 2015; Yang et al. 2017)) (Supplemental Data Table 6). Even small changes in the abundance of each protein might cumulatively contribute to the observed changes in mushroom body development or behavior in Nab2 null flies (Kelly et al. 2016; Bienkowski et al. 2017; Corgiat et al. 2022; Rounds et al. 2022). Several of the proteins encoded by transcripts with retained first introns (Trio, sick, and sif) directly regulate the activity of Rho GTPases required for axonal guidance and provide a direct link between Nab2 loss and changes in mushroom body axon morphology. Abrupt and Imp regulate the temporal specification of different subsets of mushroom body neurons (Kucherenko et al. 2012; Yang et al. 2017). For example, the transcription factor Abrupt is necessary for production of α’/β’ neurons within the mushroom body and must be downregulated in order for the later born α/β neurons to be specified (Kucherenko et al. 2012; Lin 2023). Interestingly, α’/β’ neurons are one group of neurons where both Trio and Dystroglycan (Dg) are mostly highly expressed during mushroom body development (Awasaki et al. 2000; Yatsenko et al. 2021). Thus, in addition to axonal guidance, Nab2 loss might cause changes in the temporally restricted specification of different mushroom body neurons. Future studies investigating whether Nab2 is associated directly with each of these transcripts and the ways in which Nab2 coordinates splicing, intron retention and polyadenylation will help to illuminate how this family of RNA binding proteins functions during brain development.

## Materials and Methods

### Fly stocks

All *Drosophila melanogaster* stocks were housed at 25°C on standard molasses-based food (Nutrifly MF, Genesee Scientific) according to the manufacturer’s directions, unless otherwise stated. Fly stocks containing the Nab2 null (*Nab2^ex3^*) allele and wild-type control (*Nab2^pex41^*) allele were described previously (Pak et al. 2011). All other stocks were obtained from either the Bloomington Stock Center (BSC) or the Vienna *Drosophila* Resource Center (VDRC) and are listed in Supplemental Table 7. We used FlyBase (release FB2026_02) to find information about stocks, alleles, reagents, phenotypes, interactions, and references (Ozturk-Colak et al. 2024).

### Fly eye imaging

Before using it for analysis of eye phenotypes, the *Nab2^EP3716^*allele was outcrossed to *w^1118^* (Bloomington Stock #5905) for six generations and then combined with *ninaE.GMR-GAL4* (Bloomington Stock #1104). Flies containing both *GMR-GAL4* and *Nab2^EP3716^* were crossed to flies containing a *UAS-RNAi* construct targeting a gene of interest, resulting progeny containing *ninaE.GMR-GAL4, Nab2^EP3716^*, and the specified *UAS-RNAi* construct were selected. The eyes of approximately 5-10 male and 5-10 female flies for each genotype were imaged. Only male flies are shown in Fig 1 and 2.

### Brain dissection and immunofluorescence

Wandering L3 larvae of the appropriate genotypes were collected, rinsed three times in ice-cold 1X PBS to remove attached food, and brains were dissected using Dumont #5 forceps. Brains were then fixed in 0.1% PBST (0.1% Triton-X-100 in 1X PBS) containing 4% paraformaldehyde (EMS Biosciences) for 20 minutes with rocking at room temperature. Fixative was removed, the tissue was rinsed twice with 0.1% PBST, and then the tissue was permeabilized in 0.3% PBST (0.3% Triton-X-100 in 1X PBS) for 20 minutes at room temperature with rocking. Following permeabilization, tissue was rinsed in 0.1% PBST, blocked in 5% NGS (5% normal goat serum, 0.1% Triton-X-100, 1X PBS) for 30 minutes, and then incubated for 2-3 days with primary antibodies (diluted in 5% NGS) at 4°C with rocking. Primary antibody solution was then removed, tissue was washed in 0.1% PBST, and the tissue was then incubated with secondary antibodies (diluted in 5% NGS) for 3 hours at room temperature, with rocking. Tissue was then washed in 0.1% PBST, mounted onto Superfrost Plus slides in VectaShield (Vector Labs), and imaged using an inverted Olympus Fluoview 3000 laser scanning confocal microscope. Pupal and adult brains were dissected using the same methods (Kelly et al. 2016) but without the initial PBS wash to remove excess food. Antibodies targeting Fas2 (1D4, RRID: AB_528235, (Hummel et al. 2000)) and Trio (A9.4A concentrated, RRID: AB528494, (Awasaki et al. 2000)) were obtained from the Developmental Studies Hybridoma Center (DSHB) and both diluted 1:20. Primary chicken anti-GFP antibodies were obtained from Aves Labs (GFP-1010, RRID: AB_2307313) and diluted 1:500. Alexa488-labeled goat anti-mouse secondary antibody (Invitrogen, cat A-11001, RRID: AB_2534069) was diluted 1:200.

### RNA isolation, library preparation, and RNA-seq

Brains were dissected as described above, but instead of being fixed, they were transferred to nuclease-free Eppendorf tubes and frozen at - 80°C. For each biological replicate, approximately 50-75 brains of the selected genotype were combined, homogenized in buffer RLT containing 40 mM DTT, and total RNA was isolated using Qiagen RNeasy or RNeasy Plus kits, according to the manufacturer’s directions. Genomic DNA (gDNA) was removed DNAse digestion (for RNeasy kits) or gDNA eliminator columns (for RNeasy Plus kits). RNA quality was assessed by Agilent Tape Station. Library preparation and paired-end RNA sequencing was performed by the Georgia Genomics Core at The University of Georgia, according to standard protocols. Briefly, poly(A) RNA was selected, and libraries were constructed using the KAPA Stranded mRNA-Seq kit with Capture Beads (KK8421). Samples from each biological replicate were then pooled across a single flow cell and 75-nucleotide paired-end sequencing was performed using the Illumina NextSeq platform.

For experiments utilizing RNA samples for cDNA synthesis, Nanopore sequencing, and qRT-PCR analysis, MasterPure (Lucigen/Biosearch Technologies) kits were used in place of the Qiagen RNeasy kit, according to the manufacturer’s directions. Briefly, ∼60-75 brains per genotype were dissected and frozen at -80°C until ready for use. Brains were then thawed, homogenized in Tissue and Cell Lysis buffer containing Proteinase K, and incubated for 30 minutes at 65°C to lyse cells. Proteins were precipitated by the addition of MPC precipitation buffer and lysates were centrifuged for 15 minutes at 4°C. The supernatant was removed, and nucleic acids were precipitated by the addition of isopropanol. The resulting pellet was washed in 70% ethanol, air-dried, and dissolved in nuclease-free water. DNA was then hydrolyzed by incubating RNAse-free DNAse with the samples for 30 minutes at 37°C and DNAse was precipitated with a second round of MPC precipitation buffer. RNA was then precipitated with isopropanol. Pellets were washed twice with 70% ethanol and resuspended in nuclease-free water. Finally, in order to remove contaminating salts from the MPC and lysis buffers used in the MasterPure procedure, RNA was precipitated overnight at -20°C with the addition of ethanol/sodium acetate. RNA was pelleted by centrifugation at 4°C for 15 minutes, pellets were again washed twice with 70% ethanol, and dissolved in 30µl of RNAse-free water (Qiagen).

### Nanostring nCounter gene expression array

Validation of differential gene expression was performed using a custom designed nCounter codeset. NanoString nCounter technology is based on direct detection of target molecules using color-coded molecular barcodes, providing a digital simultaneous quantification of all target RNAs. Total RNA from L3 larvae brain tissue was isolated using Rneasy Plus Mini Kits (Qiagen) as described above. Each biological replicate consisted of RNA isolated from 60-75 larval brains. 100 ng of total RNA was hybridized overnight with nCounter Reporter (8 μL) probes in hybridization buffer and in excess of nCounter Capture probes (2 μL) at 65°C for 18 h. After overnight hybridization, probes in excess were removed using two-step magnetic beads-based purification on an automated fluidic handling system (nCounter Prep Station). Biotinylated capture probe-bound samples were then immobilized and recovered on a streptavidin-coated cartridge. The abundance of specific target molecules was quantified using the nCounter digital analyzer. Individual fluorescent barcodes and target molecules present in each sample were recorded with a CCD camera by performing a high-density scan (325 fields of view). Images were processed internally into a digital format and exported as Reporter Code Count (RCC).

### Oxford Nanopore (ONT) long read sequencing

Larval brain tissue was dissected fresh as described above, pooled into batches of 15 brains, and frozen at -80°C. Once ∼60-75 brains from each genotype had been obtained, brains were thawed on ice, combined together, and RNA was isolated using Lucigen’s MasterPure Complete DNA and RNA Purification kit (now sold by Biosearch Technologies), as described above. Total RNA was then sequenced using either Oxford Nanopore’s direct RNA sequencing kit (SQK-RNA002) or cDNA sequencing kit with barcodes (SQK-PCB114.24). In order to capture full-length poly(A) tails, cDNA-RT adaptors were ligated to the 3’-end of the poly(A) tail prior to the reverse transcriptase step, as detailed in the manufacturer’s directions. For cDNA sequencing reactions, one replicate of each genotype was combined and sequenced on each flow cell.

### Illumina RNA-Seq Bioinformatics and Data Availability

Following initial quality control analyses, all reads, reference genome sequences (BDGP version 6.46), and associated gene annotations were uploaded to the servers at The Ohio Supercomputer Center (OSC). All sequencing data and analysis files have been deposited to the SRA database (BioProject ID: PRJNA1109358). All code used for bioinformatics is available as Supplemental Data File 1.

Reads were first aligned to the *Drosophila melanogaster* reference genome using STAR (ver. 2.7.9a, (Dobin et al. 2013)). The featureCounts algorithm from the Subread package (ver. 2.0.6, (Liao et al. 2014)) was then used to assemble a count matrix from each of the .bam files generated by STAR. Differentially expressed (DE) genes were then identified using DESeq2 (ver. 1.42.0, (Love et al. 2014)), using an apeglm for LFC estimates (Zhu et al. 2019). ComBat-Seq (Zhang et al. 2020a) was used initially to remove a known batch effect caused by the introduction of new RNeasy (Qiagen) reagents. Unknown batch effects were modeled using Surrogate Variable Analysis (SVA) via the svaseq function in the R/Bioconductor package sva (Leek 2014) and included in the DESeq2 design (Ritchie et al. 2015).

### Oxford Nanopore Technologies (ONT) Data Analysis

Raw signal from cDNA sequencing reactions was basecalled and demultiplexed in separate steps using Dorado (ver 1.0.0). Poly(A) tail length was also calculated during basecalling. Basecalled .bam files from Dorado were then converted to .fastq files using samtools (ver 1.21, (Li et al. 2009)) and pychopper (RRID: SCR_018966, https://github.com/nanoporetech/pychopper) was used to orient reads. Reads were then aligned to the *Drosophila melanogaster* (BDGP 6.54) genome using minimap2 (ver 2.28, (Li 2018)) and a custom python script (Supplemental File 2) was used to determine poly(A) tail lengths for all reads aligned to each gene. Due to limited sequencing depth and the length and 5’ position of retained introns in Nab2 null tissue, it was not possible to obtain accurate poly(A) tail length information for each mRNA isoform. Therefore, poly(A) tail length information was summarized at the gene level, rather than the transcript isoform level. Sequencing quality and length statistics were generated using samtools and alignment statistics were generated using SeqKit (Shen et al. 2016).

ONT direct RNA .fast5 files (from the SQK-RNA002 kit) were first converted to the .pod5 format and then basecalled using Dorado (ver 0.9.5). Reads were converted to .fastq format using samtools (ver 1.21, (Li et al. 2009)) and aligned to the *D. melanoster* genome (BDGP 6.54) using minimap2. All other steps for dRNA data analysis were identical to those performed with data from cDNA sequencing. Aligned reads were also visually inspected using IGV. All sequencing data and analysis files have been deposited to the SRA database (BioProject ID: PRJNA1109358).

### Nanostring data analysis

The NanoTube R package (Class et al. 2023) was used for analysis of Nanostring data. Raw .RCC files were imported into R Studio and the standard “nSolver” option was used to normalize the data before differential expression analysis. This option uses internal housekeeping genes (*Act5c, Act42A, RpS20, mRpL49, FoxK, eEF1alpha1,* and *eEF1delta*) and synthetic single-stranded DNA positive control probes designed by NanoString to apply a correction factor to the raw data. Internal housekeeping genes were picked based on (Lu et al. 2018). No background subtraction was performed on NanoString data. NanoTube then used limma (Ritchie et al. 2015) to identify differentially expressed genes using the housekeeping gene normalized data. Since the normalized nCounter data varied by several orders of magnitude between genes depending on expression levels and would therefore be difficult to represent on a single figure, the nCounter data was scaled using the minimum and maximum levels of expression for each gene:

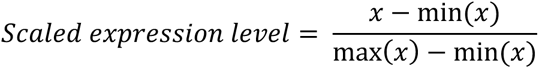

Where x is an individual data point for one gene, min(x) is the minimum level of expression in the nCounter experiment, and max(x) is the maximum level of expression for a given gene. The scaled data points for control and Nab2 null samples were used for the boxplots shown in Fig 3B, but non-scaled housekeeping-normalized expression values were used by limma for differential expression analysis.

### Analysis of intron retention by Majiq and IRFinder

Majiq (ver 2.5.1) was used according to the authors instructions (Vaquero-Garcia et al. 2023). Briefly, aligned short-read .bam files and *D. melanogaster* annotations (BDGP 6.46.110) were used by Majiq-build to create splicegraphs and define local splice variations (LSVs). When using Majiq build, defaults values were used except for –min-experiments = 0.3. Majiq deltapsi was then used to quantify the differences in abundance of LSV usage between samples. Deltapsi output was then analyzed in R. Retained intron LSVs with a probability-changing > 0.6 were further analyzed.

Intron retention ratios were also determined using IRFinder 1.3.1 (Middleton et al. 2017). IRFinder was used to build an IRFinder reference from the *Drosophila* genome (BDGP release 6.46.110). The IRFinder reference contains coordinates (i.e. nucleotide positions within the genome) for potential intron sequences between exons and omits regions of poor unique mapability, as determined by synthetic read mapping across the genome (please see (Middleton et al. 2017) for more details). This newly created intron annotation reference file was then used with name-sorted STAR .bam output files generated during differential gene expression analysis above to quantify intron retention. Intron Retention (IR) ratios from each sample were determined by IRFinder using the equation below (Middleton et al. 2017):

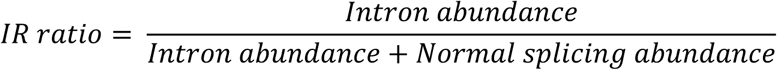

Where Intron abundance is the median intron depth and Normal splicing abundance is the number of reads connecting adjacent exons. Both intron and normal splicing abundances are normalized for overall feature length. Default IRFinder values were used (limiting minimum intron length to 50 nucleotides), and significant differences in the fold-change of intron retention between samples were then determined by using DESeq2. The coordinates of selected retained introns identified by IRFinder were visually inspected in the Integrative Genomics Viewer (version 2.13, (Robinson et al. 2011)) to determine the relative location of that intron within the entire transcript. IRFinder output was filtered to include only those introns significantly retained, p <1e^−5^ since we found that lower thresholds allowed for a higher number of introns to be marked as retained when very few reads were present by visual inspection.

### Intron length and splice-site strength determination

The length of retained introns was determined using the starting and ending coordinates of each intron identified by DESeq2 as significantly retained in Nab2 null tissue. The length of all first introns in all *Drosophila* mRNA transcripts and the length of all first introns in all canonical transcripts were determined using a custom R script and the AnnotationHub R package (ver. 3.10.0, (Morgan M 2024)). Briefly, the starting and ending coordinates for the first and second exons of each mRNA were downloaded using AnnotationHub. The ending nucleotide position of exon 1 and the starting nucleotide position of exon 2 were used to define the boundaries of the first intron. Once starting and ending coordinates were obtained, the overall length of the intron could then be calculated by subtracting the two values. The length of all introns was found in a similar manner, but not limited to the first two exons.

Splice site strength scores were determined using the Maximum Entropy model implemented into MaxEntScan (Yeo and Burge 2004). Starting and ending nucleotide coordinates of each intron were determined as described above and the DNA sequences surrounding 5’ and 3’ splice sites were obtained using the getSeq command of the BSgenome R package. For 5’ splice sites, 3 nucleotides of the exon and 6 nucleotides downstream of the splice site into the intron were obtained (9 nucleotides total). For 3’ splice sites, 20 nucleotides of the intron upstream of the 3’ splice site and 3 nucleotides downstream of the 3’ splice site in the exon were obtained (23 nucleotides total). The VarCon R package was used to call MaxEntScan and determine the splice site strength score of each 5’ and 3’ splice site from retained introns and all splice sites. Splice site scores were then compared between introns using a one-sample t-test where mu was set to the mean score of all 5’ or 3’ splice sites across the entire genome. Only introns identified as retained by both Majiq and IRFinder were used for splice site strength analyses.

### Meme Analysis of 5’ and 3’ splice sites

The genomic coordinates and sequences of 5’- and 3’- splice sites of retained introns and all first introns were obtained as detailed above. Retrieved sequences were then used for separate MEME analyses (ver 5.5.5, (Bailey and Elkan 1994; Bailey et al. 2015)) to identify common patterns in the input sequences between retained and all 5’- and 3’- splice sites. Default parameters were used except that any number of repetitions were expected (anr) and motif sites were searched on the given strand only.

### RBP binding motif enrichment analysis within retained introns

Sequences of retained introns including the surrounding upstream and downstream 50 base pairs were obtained from genomic coordinates using the getSeq command of the BSgenome R package. Retrieved sequences were then used as input for AME analysis (ver. 5.5.5, (Grant et al. 2011)) to identify over-represented sequences. The *Ray2013_rbp_Drosophila_melanogaster.dna_encoded.meme* database (provided by the MEME suite and derived from (Ray et al. 2013)) was used as a motif database. FIMO was used to identify the location of individual binding motifs within the retained intron sequences.

### Compositional analysis of adenosine content near splice sites

The frequency of adenosine around introns identified by both Majiq and IRFinder as retained was calculated using the Biostrings R package. For regions near the 5’ splice site (5’ss), 100 nucleotides (nts) into the exon upstream of the 5’ss and 50 nts downstream into the intron downstream of the 5’ss were analyzed. For regions near the 3’ splice site (3’ss), 50 nts into the intron upstream of the 3’ss and 100 nts into the downstream exon were analyzed. Sequences within each 150 nt region were obtained in R and the letterFrequencyInSlidingView function of the Biostrings package was used to compute the percentage of adenosine within a five-nucleotide sliding window.

### cDNA synthesis for qRT-PCR and semi-quantitative PCR

250 ng of total RNA from each biological replicate was reverse transcribed using Biorad’s iScript reverse transcription kit, with some modifications to the manufacturer’s directions as suggested in (Minshall and Git 2020). Total RNA was initially heated to 65°C for 10 minutes and cooled rapidly on ice. iScript reverse transcriptase and buffer (containing a mixture of oligo-d(T) and random hexamer primers) were then added to the chilled RNA, incubated at 25°C for 10 minutes and then at 46°C for an additional 30 minutes. The reverse transcriptase enzyme was then heat-inactivated by incubation of the mixture at 95°C for 1 minute. Diluted cDNA was then used as a template for both semi-quantitative PCR and quantitative real-time PCR (qRT-PCR) experiments.

### Semi-quantitative PCR and quantification of gel images

In order to confirm the presence of retained introns within specific gene isoforms, PCR reactions were performed using Taq polymerase (Qiagen) according to the manufacturer’s directions. Approximately 1ng of cDNA was used as a template in PCR reactions containing 0.3 µM forward and reverse primers. PCR products corresponding to spliced and unspliced transcripts were resolved on 1-2% agarose gels using GelRed staining and imaged using a BioRad ChemiDoc imager. Intron retention was analyzed using 5-7 biological replicates for each genotype. Band brightness was quantified in ImageJ after converting images to 8-bit files. The ratio of unspliced to spliced mRNA was determined using the following equation, where Unspliced = densitometry of the PCR product corresponding to the unspliced mRNA transcript, Spliced = densitometry of the PCR product corresponding to the spliced mRNA transcript, Background(1) = background signal near the PCR product corresponding to the unspliced transcript, Background(2) = background signal near the PCR product corresponding to the spliced transcript:

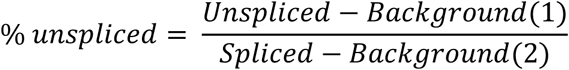

Primer sequences were designed using the PrimerQuest tool from IDTDNA.com and are included in Supplemental Data Table 8. Transcript sequences used for primer design were obtained from Ensembl Release 104 and the Berkeley *Drosophila* genome project (BDGP), version 6.32.

### Quantitative real-time PCR (qRT-PCR)

Quantitative real-time PCR was performed using SSO Advanced Universal Green Supermix (Biorad). qRT-PCR was performed in 10µl reactions with 0.2 µM forward and reverse primers on a CFX Connect (Biorad) thermocycler. Primer efficiency and optimal cDNA concentrations for qRT-PCR reactions were first determined using cDNA serial dilutions according to established protocols (Schmittgen and Livak 2008). All primers used for qRT-PCR experiments had 90-110% efficiency as suggested by (Schmittgen and Livak 2008; Fraga et al. 2014). qRT-PCR experiments were performed in triplicate on 5-7 biological replicates; each biological replicate corresponded to RNA isolated from 60-75 *D. melanogaster* L3 larval brains. For all qRT-PCR experiments, target gene expression was normalized to *Rpl32* (also called *rp49*) expression. *Rpl32* is a common housekeeping gene (Lu et al. 2018) and was chosen here as a normalization gene based on the minimal non-significant change (log2 fold change = 0.08) in *rpl32* expression observed in the Illumina RNA-seq between *Nab2^pex41^* and *Nab2^ex3^* larval brains. Threshold (C_T_) values for each sample were then averaged and the 2^−1′CT^ method (Pfaffl 2001; Schmittgen and Livak 2008) was used for relative quantification. Standard error values for the relative expression of each gene were calculated from corresponding 2^−1′CT^ values from each biological replicate. Primers used for qRT-PCR were obtained from the FlyPrimerBank (Hu et al. 2013) or designed using PrimerQuest (Integrated DNA Technologies). Primer sequences are provided in Supplemental Data Table 8.

### Immunoblots

Total protein was isolated from ∼15 larval brains or adult fly heads by homogenization of fly tissue in Radioimmunoprecipitation Assay (RIPA) buffer containing protease inhibitors (Roche). Approximately 20-30µg of total protein was separated using 4-20% SDS-PAGE (Biorad), transferred to nitrocellulose (SantaCruz) and then visualized on nitrocellulose membranes by Ponceau-S staining. Ponceau-S stained membranes were imaged and then washed extensively with dH_2_O and 1X Tris-buffer-Saline supplemented with 0.05% Tween-20 (1X TBS-T) to remove excess stain. Nitrocellulose membranes were then blocked for at least 30 minutes in 5% non-fat dry milk dissolved in 1X TBS-T (5% NFDM) and incubated overnight at 4°C in 5% NFDM with primary antibodies. Nitrocellulose was then washed extensively with 5% NFDM to remove excess primary antibody and incubated for 1 hour at room temperature with secondary antibody, diluted 1:5000 in 5% NFDM. Nitrocellulose membranes were then washed again in 1X TBS-T and incubated with Clarity ECL (Biorad) to detect bound HRP-labeled secondary antibodies. If a second primary antibody was used, blots would be washed for 2-3 hours in 1X TBS-T at room temperature, reblocked in 5% NFDM, and incubated with the additional primary antibody overnight at 4°C. Antibodies used in this study include anti-Nab2 (1:4,000, (Pak et al. 2011 2016)), anti-SmB (Y12, Invitrogen, MA5-13449, RRID: AB_10944191, 1:100), anti-trio (DSHB 9.4A (Awasaki et al. 2000), 1:500), anti-tubulin (Abcam, ab179513, 1:5,000), anti-Dg ((Deng et al. 2003), 1:10,000), and anti-Rbfox1 (also known as A2BP1, 1:5,000, (Tastan et al. 2010)), HRP-labeled goat-anti mouse secondary antibody (Santa Cruz, sc-2055, 1:5,000), HRP-labeled goat-anti rabbit secondary antibody (Santa Cruz, sc-2004; 1:5,000), and HRP-labeled goat anti-guinea pig (ThermoFisher, A18769, RRID: AB_2535546, 1:5,000).

### Generative AI usage statement

Generative AI models (Claude: Sonnet 5 and ChatGPT5.5) were utilized to help analyze Oxford Nanopore dRNA and cDNA sequencing results, to formulate code for the sliding window analysis used in Figure 7, and to generate code for statistical tests in R. Generally, one of the AI models (usually Sonnet 5) was used to generate code for these analyses and then the other AI model (usually ChatGPT5.5) was used to double check code for errors. No AI models were used in the writing or editing of the manuscript text.

## Acknowledgements

We would like to acknowledge the generous research sabbatical/leave program, Luce Foundation funds, and Faculty Development funds provided by The College of Wooster to S.M.K. that allowed for much of this work and writing to be accomplished. We would also like to thank Ken Moberg and Jacob Kagey for critical reading of manuscript drafts. Stocks obtained from the Bloomington *Drosophila* Stock Center (BDSC, NIH P40OD018537) and Vienna *Drosophila* Resource Center (VDRC (Dietzl et al. 2007)) were used in this study. Illumina and dRNA ONT sequencing were performed by the Georgia Genomics and Bioinformatics Core at The University of Georgia (GGBC, UG Athens, GA, RRID:SCR_010994). cDNA ONT sequencing was performed by The Ohio State University College of Food, Agricultural, and Environmental Sciences (CFAES, Wooster, OH). Bioinformatics analysis of sequencing information was performed using resources from The Ohio Supercomputer Center (OSC, https://ror.org/01apna436) located at The Ohio State University. We thank the Genomics Shared Resource at The Ohio State University Comprehensive Cancer Center (OSU CCC), Columbus, OH for conducting the NanoString/nCounter analyses. We would also like to thank Michael Buszczak and Wu-Min Deng for graciously sharing anti-Rbfox1 (A2BP1) and anti-Dg antibodies, respectively.

