## Supplementary material for "The polyadenosine RNA binding protein Nab2 regulates alternative splicing and intron retention during *Drosophila melanogaster* brain development": Supp Fig Legends

Supplemental Figure Legends

**Supplemental Fig 1: A small number of reads map to the Nab2 locus in RNA samples isolated from Nab2 null larval brain tissue.** RNA was isolated from both wild-type (genotype: *Nab2^pex41^/Nab2^pex41^*) and Nab2 null (genotype: *Nab2^ex3^/Nab2^ex3^*) larval brain tissue and used for Illumina short-read sequencing. Reads were mapped to the *Drosophila* genome and visualized using the Integrative Genome Viewer (IGV). (A) IGV screenshot of the Nab2 genomic locus in four wild-type samples and four Nab2 null samples. Very few reads were observed at the Nab2 locus in Nab2 null samples. All tracks are scaled to the same level. (B) Principal Component Analysis (PCA) plots of RNA-seq samples following differential gene expression analysis by DESeq2 after removal of batch effects. Wild-type control ($Nab2^{pex41}/Nab2^{pex41}$, n = 6) samples cluster independently of Nab2 null ($Nab2^{ex3}/Nab2^{ex3}$, n = 7) samples along principal component 1 (PC1).

**Supplemental Fig 2: Nab2 loss alters the expression of genes involved in specific molecular functions.** Enriched Gene Ontology (GO) terms associated with differentially expressed genes (adjusted p-value <0.05, $\log_2$-fold change < -1 or >1) in Nab2 null ($Nab2^{ex3}/Nab2^{ex3}$) larval brains were identified by clusterProfiler. Nab2 loss preferentially alters genes encoding proteins with monooxygenase activity, enzymes that bind heme/iron, enzymes implicated in steroid hormone biosynthesis, and fatty-acid-CoA ligases.

**Supplemental Fig 3: Genes expressing transcripts with retained first introns in Nab2 null larval brains are also more highly expressed.** (A) All expressed genes in Nab2 null larval brains were divided into two groups: Nab2-independent genes (green, n = 10,625), which produce transcripts that do not contain a retained intron in Nab2 null larval brains and Nab2-dependent genes (purple, n = 91), which produce transcripts containing a retained intron in Nab2 null larval brains. Nab2-dependent genes that produced intron-containing transcripts were significantly more likely to be differentially expressed (adjusted p-value as determined by DESeq2 < 0.05) between control and Nab2 null larval brains (p < 0.001, Fisher's Exact Test, Odds-ratio: 4.605, 95% CI: 2.97-7.13). (B) Nab2 is required for the correct expression of specific *Trio* and *Rbfox1* isoforms. Many isoforms of other Nab2-dependent genes that produce intron-containing transcripts are also slightly increased in Nab2 null larval brains. RNA was isolated from control ($Nab2^{pex41}/Nab2^{pex41}$) and Nab2 null ($Nab2^{ex3}/Nab2^{ex3}$) larval brain tissue, reversed transcribed, and resulting cDNA was used as a template for RT-qPCR reactions with gene specific primers that anneal to exons surrounding non-retained introns. Obtained $C_T$ values for target genes were normalized to $C_T$ values for the housekeeping gene *rp49* (also called *rpl32*) using the $2^{-\Delta CT}$ method to obtain normalized expression values. For plotting purposes only, normalized expression values were scaled so that the minimum expression value from replicates for each transcript was set to 0 and the maximum expression value from replicates for each transcript was set to 1. Boxplots show these scaled rp49-normalized expression values. Normalized but non-scaled $2^{-\Delta CT}$ values were used for statistical analyses. *Trio* - isoform D (RD) and select *Rbfox1* isoforms (EHIKLM) are significantly elevated in Nab2 null brains. Isoforms containing retained introns are not significantly altered between control and Nab2 null larval brains (* = p<0.05, Wilcoxon test, used with Benjamini-Hochberg corrections).

**Supplemental Fig 4: Nab2-dependent genes that produce intron-containing transcripts in Nab2 null larval brains are involved in developmental, behavioral, and cell signaling biological processes.** Enriched gene ontology (GO) terms associated with genes identified by both Majiq and IRFinder as producing transcripts with a retained intron in Nab2 null larval brains ("Nab2-dependent genes") were identified by clusterProfiler. (A) Nab2-dependent genes were enriched for biological functions such as morphogenesis (GO: 0007560), neurotransmitter secretion and synapse function (GO: 0046928, 0007416, and 0050803), axon guidance and cell adhesion, (GO: 0007411, 0007155, 0050839), and behavior (GO: 0007610 and 0007619). Several cellular compartments, including cell junction (GO: 0030054), cell cortex (GO: 0099738), and axons (GO: 0030424), were also enriched in Nab2-dependent genes. Data points for each GO term are plotted according to its Benjamini-Hochberg adjusted p-value and sized according to its fold enrichment. The fold-enrichment is calculated as explained (Yu et al. 2012). See Supplemental Table 5 for all significantly enriched GO terms and genes included in each term. (B) The genes for the top seven mostly highly enriched biological functions were plotted using the cnetplot R package to show interconnectedness between terms.

**Supplemental Fig 5: Direct RNA (dRNA) sequencing reveals that transcripts containing retained first introns in Nab2 null larval brains are otherwise correctly spliced.** (A) Oxford Nanopore Technologies (ONT) direct RNA sequencing of total RNA samples isolated from control or Nab2 null *Drosophila melanogaster* larval brain tissue demonstrated that very few transcripts are produced in Nab2 null larvae. Reads that aligned with the *Nab2* genomic locus were visualized with the Integrative Genomic Viewer (IGV) software. Blue bars at the top show the *Nab2* gene and other nearby genes. Black arrows and blue triangular markers indicate the direction and template strand of transcription. Thin blue boxes represent non-coding exons while thicker blue boxes represent coding exons. Thin lines between exons represent intronic sequences. In wildtype flies ($Nab2^{pex41}/Nab2^{pex41}$), aligned ONT reads cover the entire *Nab2* locus and are spliced correctly. In RNA from Nab2 null ($Nab2^{ex3}/Nab2^{ex3}$) larval brain tissue, a small number of reads aligned with the *Nab2* locus but many of these originate from an upstream noncoding RNA (*CR46093*). The region of the Nab2 locus deleted by imprecise excision of *EP3617* (Pak et al. 2011) are shown by dotted lines. (B and C) IGV tracks at the *AP-1*$\gamma$ (B) and *Rbfox1* (C) loci are shown as examples of highly expressed and more moderately expressed genes, respectively. While transcripts from wild-type larval brain tissue mainly contain correctly spliced first introns, the sequenced transcripts produced in Nab2 null larval brains have retained first introns. RNA from both wild-type and Nab2 null samples are correctly spliced at more downstream exon-exon junctions.

**Supplemental Fig 6: The binding motifs of multiple RNA Binding Proteins are enriched in and near introns retained in Nab2 null larval brains.** (A) The sequence of each retained intron (identified by both Majiq and IRFinder) in Nab2 null larval brains and the upstream and downstream 50 nucleotides (nts) into surrounding exons were used by FIMO (Grant et al. 2011) to identify the presence of RNA binding protein (RBP) binding motifs. Arrest (Aret, also called Bruno1), sex-lethal (Sxl), papi, and smooth (sm) binding sequences were the most abundant within retained intron sequences. A2BP1 (also called Rbfox1, highlighted in orange) sequences were present at low levels within retained intron sequences, compared to other RBPs. (B) Binding sequences for many RBPs, including Aret/Bru1, Sm, and Papi, are significantly enriched (Fisher's exact test, adjusted p-value $< 0.05$) in and around retained intron sequences. The sequence of retained introns and upstream/downstream 50 nts was analyzed by AME (McLeay and Bailey 2010) to identify significantly enriched RBP binding motifs.

**Supplemental Fig 7: Nab2 null larval brain tissue preferentially expresses transcripts encoding *Dystroglycan* (*Dg*) isoform C**. (A) PCR primers used to verify changes in *Dg* isoform expression anneal in exons 8 and 11. Since only *Dg* isoforms C and D contain exon 8, primers SMK489 and SMK490 should amplify only those isoforms. (B) Nab2 null brain tissue preferentially expresses *Dg*-isoform C and not *Dg*-isoform D. RNA was isolated from Nab2 null larval brain tissue or wild-type larval brain tissue and used for cDNA synthesis. Diluted cDNA was then used for semi-quantitative PCR with *Dg*-specific primers SMK489 and SMK490. PCR products were resolved using a 1% agarose gel and visualized with GelRed. Due to the absence of exon 9, PCR products derived from *Dg* isoform C should be slightly smaller in size than those obtained from amplification of *Dg* isoform D. Control samples show two PCR products, corresponding to Dg isoforms C and D, while samples from Nab2 null larval brains mainly have one, smaller PCR product which corresponds to *Dg* isoform C.

Grant CE, Bailey TL, Noble WS. 2011. FIMO: scanning for occurrences of a given motif. *Bioinformatics* **27**: 1017–1018.

McLeay RC, Bailey TL. 2010. Motif Enrichment Analysis: a unified framework and an evaluation on ChIP data. *BMC Bioinformatics* **11**: 165.

Pak C, Garshasbi M, Kahrizi K, Gross C, Apponi LH, Noto JJ, Kelly SM, Leung SW, Tzschach A, Behjati F et al. 2011. Mutation of the conserved polyadenosine RNA binding protein, ZC3H14/dNab2, impairs neural function in Drosophila and humans. *Proc Natl Acad Sci U S A* **108**: 12390–12395.

Yu G, Wang LG, Han Y, He QY. 2012. clusterProfiler: an R package for comparing biological themes among gene clusters. *OMICS* **16**: 284–287.
