## Supplementary figures and images for "The polyadenosine RNA binding protein Nab2 regulates alternative splicing and intron retention during *Drosophila melanogaster* brain development"

### Supp Fig 1

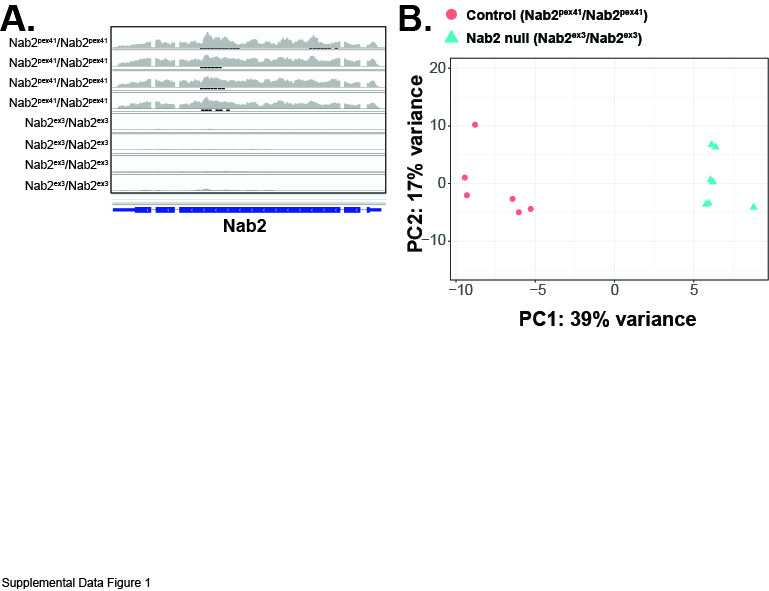

### Supp Fig 2

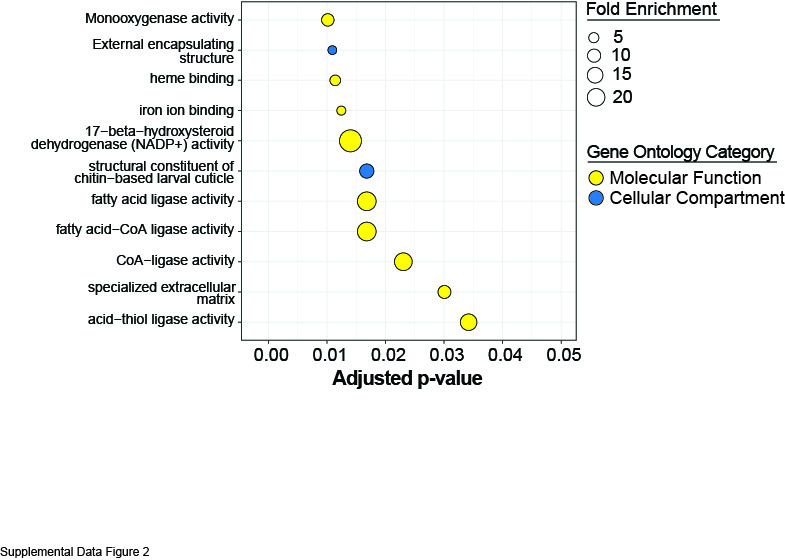

### Supp Fig 3

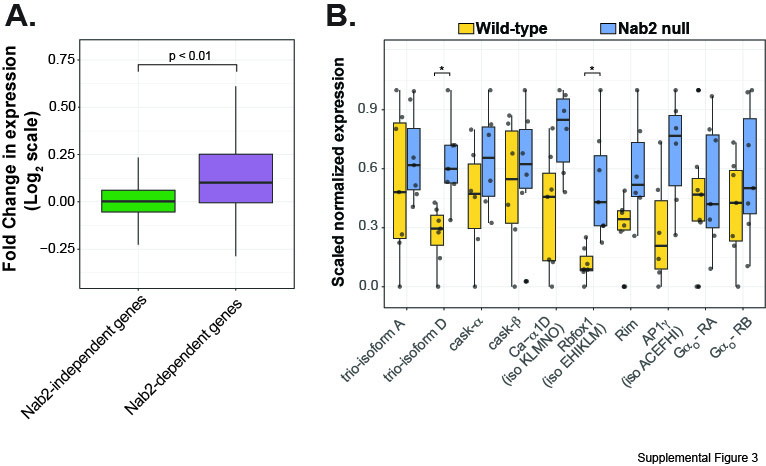

### Supp Fig 4

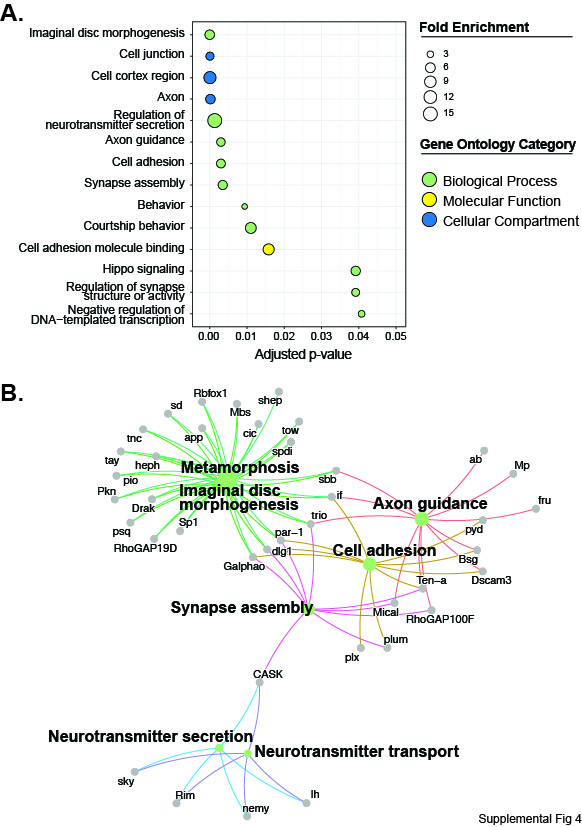

### Supp Fig 5

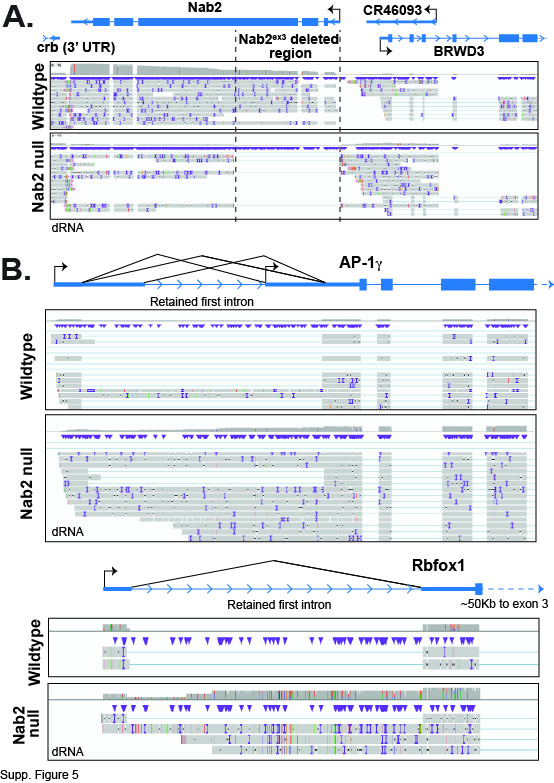

### Supp Fig 6

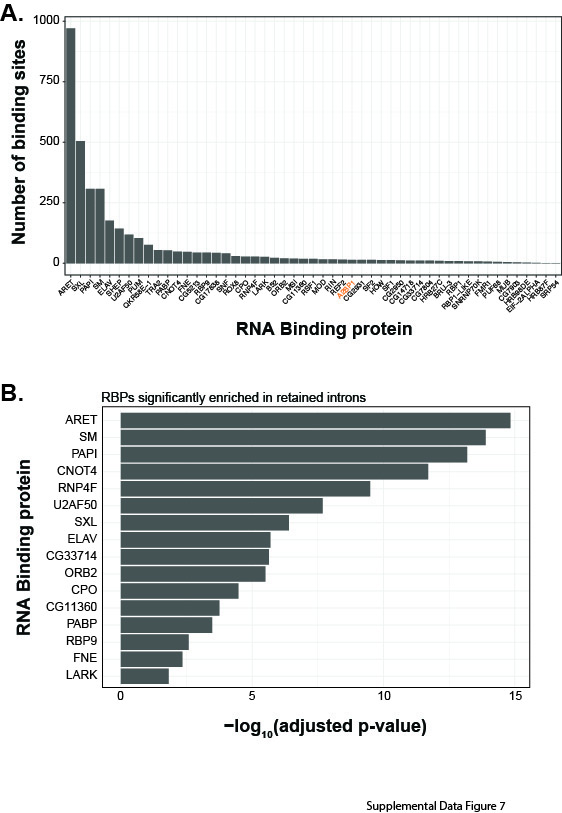

### Supp Fig 7

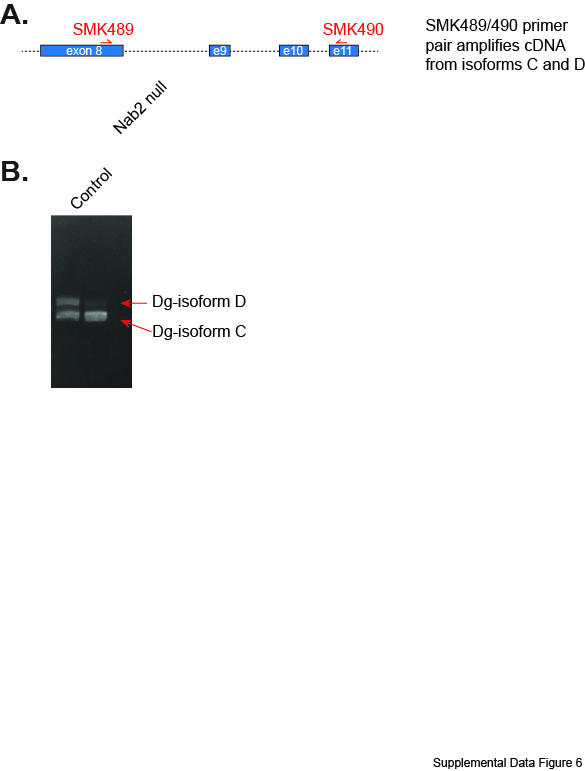
